# CNVPipe: An enhanced pipeline for accurate analysis of copy number variation from whole-genome sequencing

**DOI:** 10.1101/2025.05.18.654763

**Authors:** Jiahong Sun, Nonthaphat Kent Wong, Zhiwei Jiang, Ho-Ming Luk, Fai Man Lo, Da Di, Shijing Zhang, Raymond CB Wong, Wenfei Jin, Shea Ping Yip, Chien-Ling Huang

## Abstract

Copy number variations (CNVs) are critical contributors to the genetic architecture of complex diseases, yet many existing pipelines for whole-genome sequencing (WGS) data exhibit persistently high false discovery rates (FDR). Here, we introduce CNVPipe, an enhanced workflow that integrates widely used CNV-calling tools with a novel machine-learning framework to achieve lower FDR and higher sensitivity. CNVPipe also provides a specialised module for single-cell CNV analysis, featuring a new model that accurately estimates the ploidy of single cells to get integer copy number profile. Benchmark evaluations demonstrate that CNVPipe outperforms current pipelines in various genomic contexts. In addition to detecting recurrent CNVs in paediatric developmental disorders, CNVPipe enables large-scale functional genomics applications involving stem cell technologies. Moreover, its application to sparse single-cell DNA sequencing data provides a new aspect of research in cancer evolution. Collectively, these findings underscore the versatility and reliability of CNVPipe as a comprehensive solution for CNV detection in WGS-based research.

## Introduction

Copy number variations (CNVs), a subtype of structural variations (SVs)^1^, consist of genomic segments that differ in copy number among individuals. These alterations have been shown to influence tumour development and progression by modifying gene dosage and expression, thus contributing to cancer complexity^2^. In addition, CNVs have been linked to neuropsychiatric disorders such as autism spectrum disorders^3^ and schizophrenia^4^, where duplications or deletions of specific regions can disrupt gene function and lead to pathological phenotypes. Whole-genome sequencing (WGS) has emerged as a powerful tool for identifying these CNVs, surpassing conventional cytogenetic and array-based techniques by offering higher resolution and an extensive genome-wide scope^5^.

Numerous bioinformatics methods have been introduced for CNV detection from short-read WGS data. These methods incorporate diverse strategies such as read-depth, split-read, read-pair, and *de novo* assembly. For instance, cn.MOPS^6^, ERDS^7^, CNVKit^8^, and CNVpytor^9^ rely on assessing read depth across the genome, whereas Pindel^10^ capitalises on split-read mapping. Paired-end read approaches such as PEMer^11^ and BreakDancer^12^ focus on deviations from expected insert sizes, while Lumpy^13^, Delly^14^, and Manta^15^ use both split-read and read-pair data. Although these tools have advanced CNV detection capabilities, high false discovery rates (FDR) continue to pose a significant challenge^16,17^.

To address elevated FDR, ensemble methods integrate outputs from multiple CNV callers, as demonstrated by MetaSV^18^, sv-callers^19^, clinSV^20^, and Parliament2^21^. While these approaches often reduce FDR, merging the results from several tools can compromise sensitivity if too strict, or inflate FDR if too lenient. Additionally, variations in sequencing depth can impact the effectiveness of specific callers, emphasising the necessity for flexible strategies that adapt to different depth levels. Many existing ensemble pipelines also overlook essential steps such as upstream read processing and post-calling prioritisation, annotation, and pathogenicity assessment.

Machine learning algorithms are increasingly adopted to address these limitations in CNV detection. CN-learn^22^ incorporates multiple algorithms with random forest (RF) classifiers for exome sequencing, and EnsembleCNV^23^ combines various machine learning models for SNP array data. DudeML^24^ utilises an RF classifier trained on read depth signals, highlighting the advantages of machine learning methods for refining CNV analysis.

Against this backdrop, we developed CNVPipe, a novel ensemble CNV calling workflow encompassing all key stages: read pre-processing, parallel calling, merging, and filtering of CNVs alongside pathogenicity prediction and result visualisation. Built within the Snakemake framework^25^, CNVPipe is highly user-friendly, employs a flexible merging strategy calibrated to different sequencing depths, and introduces innovative scoring metrics to refine CNV calls. Crucially, we incorporate a support vector machine classifier to distinguish reliable CNVs from potential false positives, and this results in robust performance on both simulated and real-world benchmark datasets.

CNVPipe exhibits broad utility across diverse biological and clinical fields where CNVs serve as pivotal markers. For example, in stem cell research, it helps monitor genomic stability^26^; in genome editing experiments, it flags unintended deletions or duplications near CRISPR/Cas9 target sites^27^; and in cancer research, it detects tumourigenic CNVs and facilitates clonal evolution analyses^28^. Furthermore, CNVPipe sheds light on pathogenic CNVs in developmental and neuropsychiatric conditions^29^, and this demonstrates its potential to significantly advance research and clinical diagnostics through more precise CNV identification.

## Results

### CNVPipe workflow

CNVPipe is a unified computational pipeline built on the Snakemake framework (Github: https://github.com/sunjh22/CNVPipe), designed to streamline and automate the entire process of CNV analysis (Fig. 1). It encompasses multiple stages, beginning with raw read pre-processing or optional BAM input, followed by CNV calling, merging, scoring, and visualization. In the pre-processing step, CNVPipe handles essential tasks such as removing adaptor contamination, eliminating low-quality reads, and aligning the remaining reads to a reference genome. Recognising that resolution can significantly affect detection accuracy, we established a method to determine the optimal bin size based on sequencing depth.

**Fig. 1.**
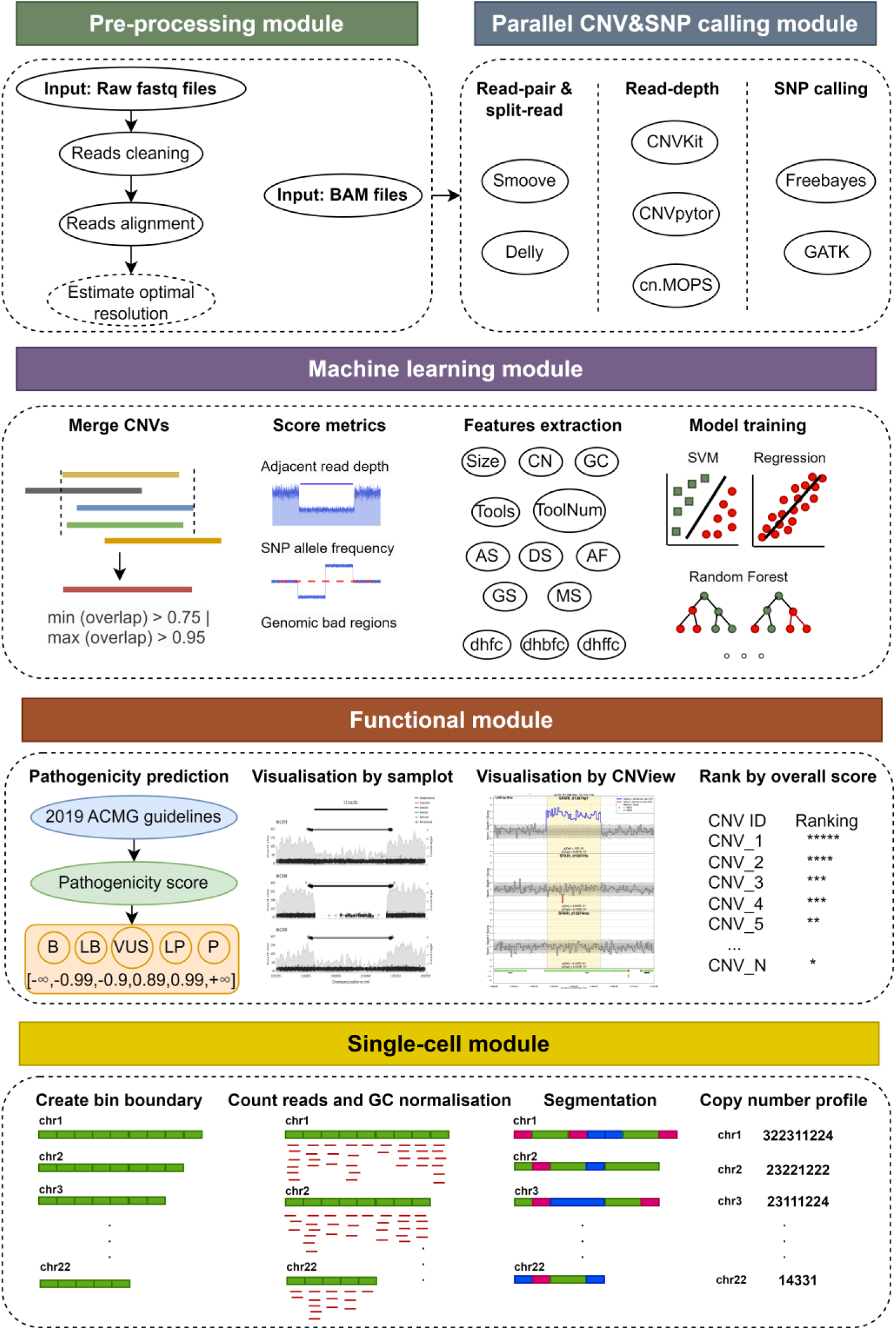
Schematic presentation of CNVPipe. CNVPipe includes five main modules: pre-processing module for reads quality control and alignment; parallel CNV calling module for detecting CNVs by five tools; machine-learning module for merging and scoring CNVs, extracting features from CNVs and training classifiers to filter CNVs; functional module for pathogenicity prediction, visualisation confirmation and overall ranking; single-cell module for detecting CNVs from sparse single-cell whole-genome sequencing data.

Within the CNV calling module, CNVPipe integrates five established CNV tools - Smoove, Delly, CNVKit, CNVpytor, and cn.MOPS, each capitalising on distinct signals (e.g., read-depth, split-read, read-pair). For samples with more than 5× coverage, CNVPipe also conducts SNP calling (e.g., via Freebayes or GATK) to refine and correct CNV calls using B-allele frequency. A novel, depth-aware merging strategy then combines the CNVs reported by these tools: low-depth data (≤ 5×) emphasises read-depth methods, whereas high-depth data (> 5×) gives priority to split-read and read-pair approaches.

To distinguish genuine from spurious CNVs, CNVPipe calculates multiple scoring metrics reflecting adjacent read-depth patterns, SNP allele frequency, overlap with repetitive or low-complexity regions, CNV size, and GC content. These metrics feed into a machine learning classifier, which can optionally be trained on a user-provided reference set or simulated data, to filter out false positives effectively. Following classification, CNVPipe compiles a comprehensive report that includes refined CNV calls, coverage and B-allele frequency (BAF) plots, and a summary of supporting evidence. In addition to analysing bulk WGS data, CNVPipe provides a specialised module for sparse single-cell WGS, enabling the construction of integer copy number profiles by binning sparse reads, applying GC normalisation, merging consecutive bins, estimating ploidies, and generating a single-cell copy number matrix for downstream analyses (Supplementary Fig. 1).

### Performance evaluation of CNVPipe

To systematically assess the performance of CNVPipe, we simulated 24 datasets spanning four CNV size ranges (10 kb, 50 kb, 200 kb, and 1 Mb) and six sequencing depths (0.1×, 0.5×, 1×, 5×, 10×, 30×), each with six replicates (Fig. 2a). Across 11 CNV callers evaluated, detection rates generally improved with increased depth, particularly for larger CNVs (200 kb to 1 Mb). However, read-depth-based methods underperformed at high coverage when identifying small CNVs, whereas split-read and read-pair tools excelled in precisely mapping smaller events at higher depths but failed at extremely low coverage (e.g., 0.1×) (Supplementary Fig. 2).

**Fig. 2.**
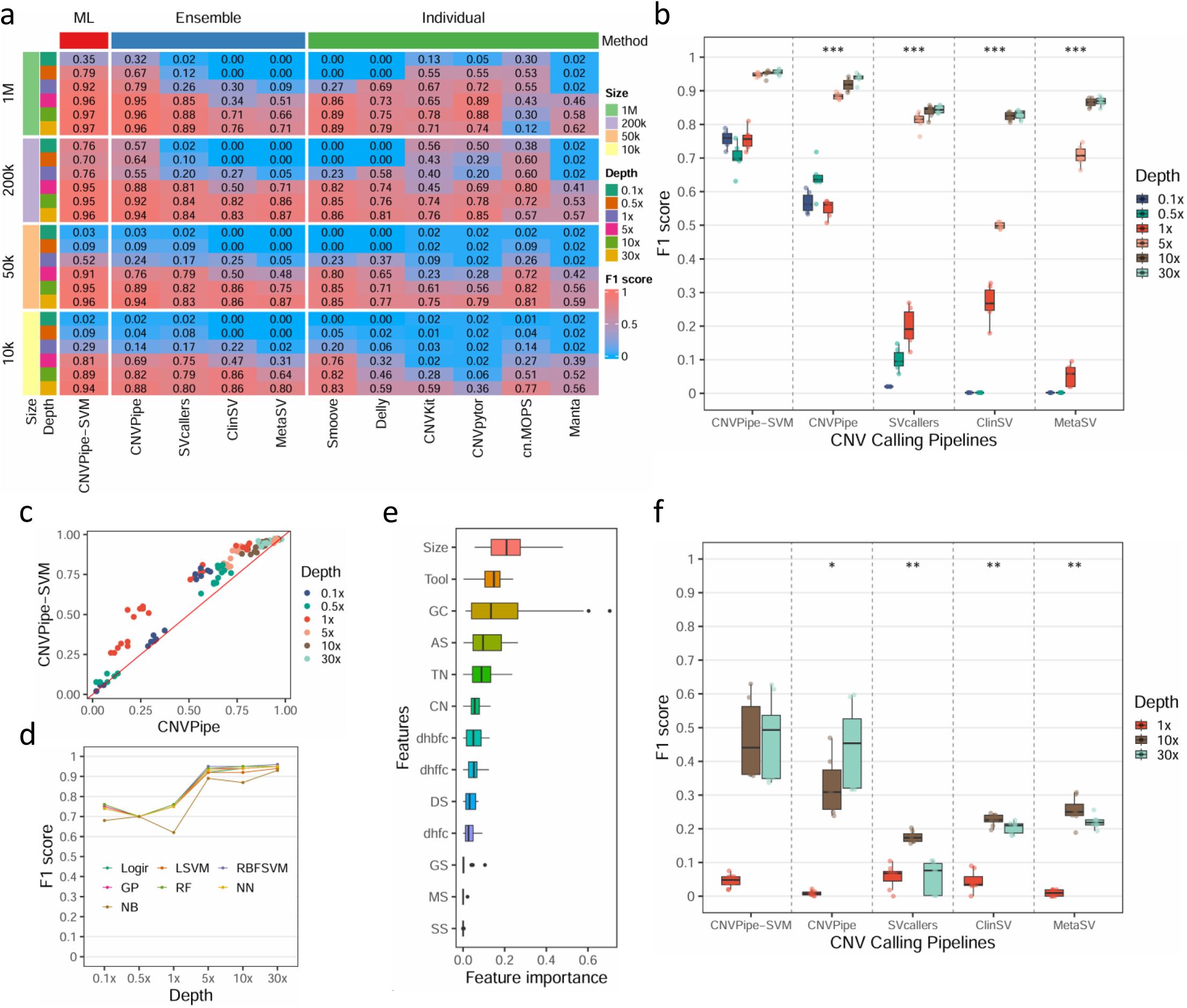
Performance evaluation of CNVPipe and other ensemble CNV callers. **a** Heatmap showing the F1 score of CNV callers under different sequencing depths and CNV sizes for simulated datasets. Column represents CNV callers, row represents different conditions (4 CNV sizes by 6 sequencing depths), numbers in each cell indicate F1 scores. ML: machine learning, SVM: support vector machine. **b** Boxplot showing the comparison of performance among CNV callers for simulated datasets (CNV size is 200 kb). Significance level is labelled at the top of boxplot. **c** Dotplot showing F1 score of CNVPipe-SVM and CNVPipe for all simulated samples. **d** Performance of different machine learning classifiers for simulated samples. Logir: logistic regression; LSVM: linear support vector machine; RBFSVM: radial basis function support vector machine; GP: Gaussian process; RF: random forest; NN: neural network; NB: naïve Bayes. **e** Discriminatory power of 13 features in support vector machine classifiers. **f** Boxplot showing the comparison of performance among CNV callers for authentic datasets. Sequencing depth 1×, 10× and 30× are included. Wilcoxon test *: *p* < 0.05, **: *p* < 0.01, ***: *p* < 0.001.

By blending these complementary strategies, ensemble pipelines outperformed individual methods. CNVPipe consistently showed higher sensitivity and reduced FDR across all size ranges and depths, and the machine learning-based CNVPipe-SVM emerged as the top performer in minimising spurious calls (Fig. 2b, c, and Supplementary Fig. 3). To further explore how different classifiers influence detection, we incorporated 13 CNV-related features - encompassing CNV size, coverage patterns, and tool-based metrics, and trained seven algorithms, including Support Vector Machine (SVM), Naïve Bayes (NB), and Random Forest (RF). On simulated datasets, most classifiers delivered comparable performance with radial basis function support vector machine (RBFSVM) and RF exhibiting marginal gains, while NB lagged behind for smaller and larger CNVs (10 kb and 1 Mb) (Fig. 2d and Supplementary Fig. 4). For RBFSVM classifier, permutation-based feature importance indicated five prominent factors (CNV size [Size], tool type [Tool], GC content [GC], accumulative score [AS], and number of tools [TN]) strongly influenced classification (Fig. 2e), and these factors shifted in importance under different sequencing depths (Supplementary Fig. 5).

When tested on real WGS data with known structural variants/CNVs, CNVPipe and CNVPipe-SVM consistently exhibited lower false discovery rates (< 0.35) than other tools, leading to superior F1 scores (Fig. 2f and Supplementary Fig. 6). Although ClinSV achieved higher sensitivity, its FDR exceeded 0.85 (Supplementary Fig. 6), reducing its practical utility for precise CNV detection. In contrast, the balanced combination of high precision and moderate recall positions the CNVPipe, along with CNVPipe-SVM, as a more reliable option for clinical and research scenarios that demand minimal false positives. A final round of comparisons involving distinct machine learning classifiers on empirical data reaffirmed RBFSVM as the leading approach for detection and F1 score (Supplementary Fig. 7). Permutation analysis discovered four features (in the order of Tool, Size, AS and GC) playing critical roles in CNV classification on real WGS data (Supplementary Fig. 8).

### Experimental validation of CNVPipe for monitoring genome stability in iPSCs and evaluating CRISPR/Cas9 off-target effects

To investigate whether CNVPipe can detect subtle genomic alterations during extended culture, we analysed CNVs of induced pluripotent stem cells (iPSCs) at passage 5 (P5) and passage 30 (P30) (Fig. 3a). CNVPipe-SVM identified 225 and 209 CNVs in P30 and P5, respectively, in which 173 CNVs (66.3%) are common among them (Fig 3b). Sv-callers – the second best ensemble CNV-calling tool in our benchmark, identified 761 and 730 CNVs in P30 and P5, in which 614 CNVs (70%) are common (Supplementary Fig. 9). Therefore, the CNV consistency rate from CNVPipe-SVM and sv-callers are comparable. We used CNView^30^ to manually inspect all the CNVs, and observed higher true positive rate for CNVPipe-SVM (65.1% and 61.3% for P5 and P30) compared with sv-callers (42.3% and 40.9% for P5 and P30) (Fig. 3c). Notably, CNVPipe-SVM identified multiple unique CNVs that failed to be detected by sv-callers (Supplementary Table 1). Subsequent droplet digital PCR (ddPCR) assays validated most of these CNVPipe-exclusive CNVs (Fig. 3d and Supplementary Table 2), attesting to the pipeline’s high sensitivity and its suitability for tracking genomic integrity throughout prolonged iPSC expansion.

**Fig. 3.**
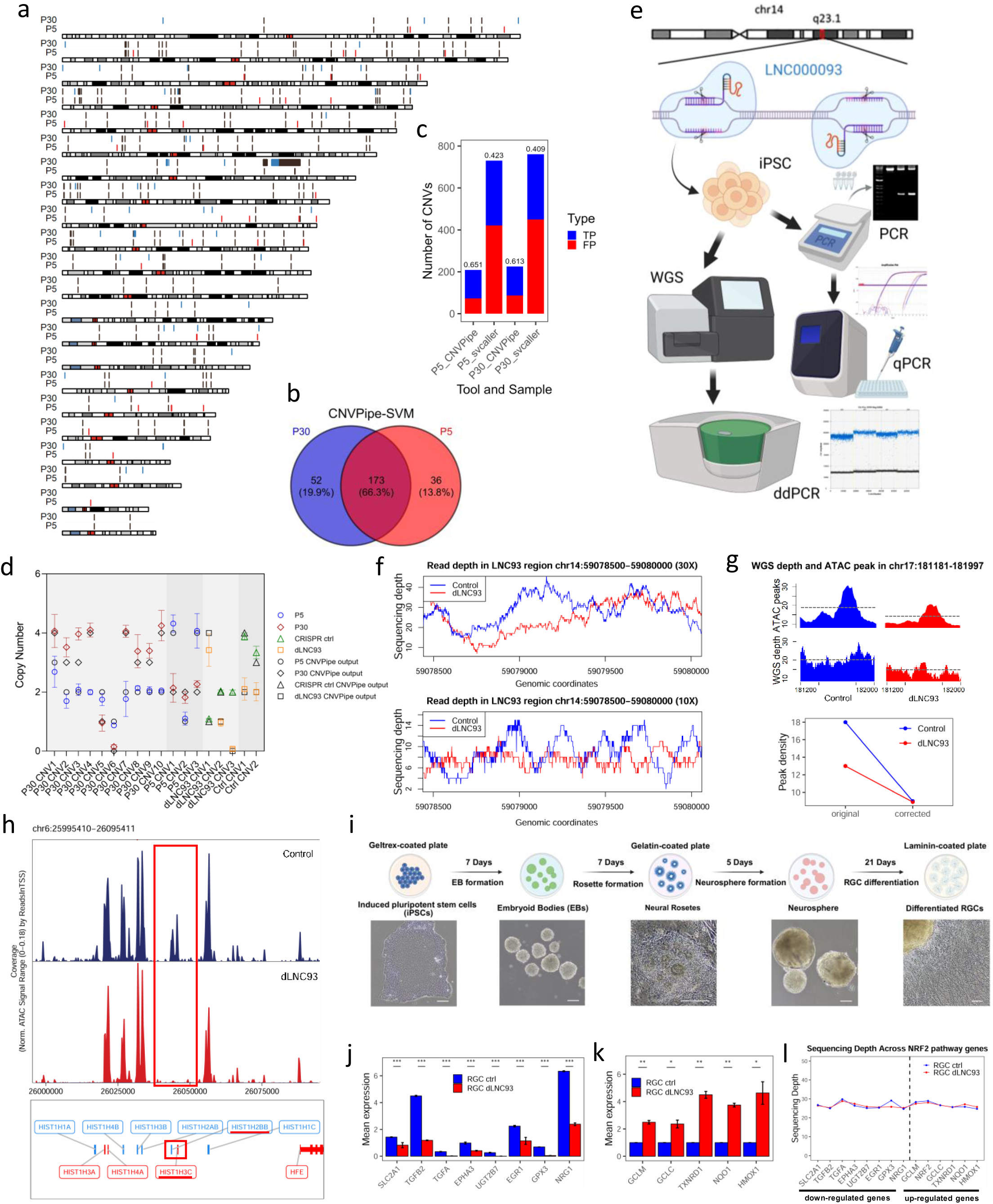
Experimental validation of CNVPipe for monitoring genome stability in iPSCs and evaluating CRISPR/Cas9 off-target events. **a** Karyoplot showing genomic position of CNVs identified in P30 and P5, green represents P30 unique CNVs, red represents P5 unique CNVs, and brown represents common CNVs between P30 and P5. **b** Percentage of unique and common CNVs in P30 and P5. **c** Percentage of true positive CNVs from CNVPipe-SVM and sv-callers manually inspected by CNView. TP: true positive, FP: false positive. **d** Droplet digital PCR (ddPCR) assays confirm most of CNVPipe-exclusive variants. **e** Workflow of CRISPR deletion of LNC000093 (dLNC93), whole-genome sequencing, quantitative PCR and droplet digital PCR. **f** Read depth of dLNC93 and CRISPR control samples in LNC93 region from 30x and 10x data. **g** ATAC peak density and WGS coverage depth in genomic region chr17:181181-181197, whose low peak value in dLNC93 sample might result from copy number deletion in this region. **h** ATAC peaks showing down-regulation of *HIST1H3C* and *HIST1H2BB* genes in dLNC93 samples. **i** Workflow of differentiating iPSCs into retinal ganglion cells. **j** Post-deletion differentiated retinal ganglion cells (RGCs) exhibit elevated expression of antioxidant genes (e.g., *NQO1*, *GCLC*, *GCLM*, *HMOX1*, and *TXNRD1*). **k** Reduced expression of genes related to NRF2 pathway (e.g., *SLC2A1*, *TGFB2*, *TGFA*, *EPHA3*, *UGT2B7*, *EGR1*, *GPX3*, *NRG1*). **l** Comparison of sequencing depth between dLNC93 and control samples on 14 genes related to NRF2 pathway. *: *p* < 0.05, **: *p* < 0.01.

We next explored the performance of CNVPipe in assessing off-target genomic alterations following CRISPR/Cas9 editing (Fig. 3e). In iPSC samples before and after the deletion of LNC000093 (control and dLNC93), we observed a pronounced drop in read coverage at the targeted locus (Fig. 3f), indicating the successful deletion of targeted region. Using CNVPipe-SVM and sv-caller to analyze control and dLNC93 samples, we observed a CNV consistency rate of 52.5% and 71.4%, respectively (Supplementary Fig. 10). The manual inspection of CNVs by CNView revealed that CNVPipe reached 96.4% and 97.6% true positive rate in control and dLNC93 samples, which is greatly higher than 58.1% and 58.5% of sv-caller, reaffirming the superiority of CNVPipe-SVM (Supplementary Fig. 10c). CNVPipe-SVM identified 4 additional CNVs that are uniquely presented in control or dLNC93 samples, which are not discovered by sv-caller, and further confirmed by ddPCR (Fig. 3d).

To test whether hidden CNVs confound downstream epigenomic and transcriptomic assays, we performed ATAC-seq and RNA-seq on control and dLNC93 iPSCs after embryoid-body induction. CNVPipe flagged a single CNV (chr17:60,000-255,592) that overlapped one of 299 down-regulated ATAC peaks (chr17:181,181-181,997). Removal of this CNV-contaminated region markedly improved peak normalisation, confirming that copy-number dosage—not chromatin remodelling—explained the apparent accessibility loss (Fig. 3g). Gene-score analysis of the corrected ATAC profiles revealed concordant down-regulation of two histone-variant genes, *HIST1H3C* and *HIST1H2BB*, in both dLNC93 iPSCs and embryoid bodies (Fig. 3h), implicating LNC000093 in chromatin regulation. Importantly, CNVPipe verified that neither gene resides within a CNV, ruling out dosage artefacts.

The same iPSC lines were then differentiated into retinal ganglion cells (RGCs) following a standard 40-day protocol (Fig. 3i). Despite normal morphogenesis, dLNC93-derived RGCs showed a transcriptional shift consistent with NRF2 activation—up-regulation of antioxidant genes (*GCLM*, *GCLC*, *TXNRD1*, *NQO1*, *HMOX1*) and suppression of stress-sensitive loci (*SLC2A1*, *TGFB2*, *TGFA*, *EPHA3*, *UGT2B7*, *EGR1*, *GPX3*, *NRG1*) (Fig. 3j, 3k and Supplementary Fig. 11). CNVPipe again confirmed that none of these NRF2-pathway genes carried inadvertent deletions or amplifications (Fig. 3l and Supplementary Fig. 12), demonstrating the value of the pipeline for ruling out off-target copy-number artefacts in functional genomics experiments.

Together, these results illustrate how CNVPipe not only improves CNV detection accuracy but also safeguards multi-omic read-outs, enabling confident interpretation of gene-editing studies in stem-cell models.

### Identification of potential pathogenic and regulatory CNVs in paediatric patients with developmental defects using CNVPipe

We applied CNVPipe to 150 WGS datasets from paediatric patients presenting various developmental conditions, such as developmental delay (DD), autism spectrum disorder (ASD), and multiple congenital anomalies (MCA) (Supplementary Fig. 13). A heatmap illustrates each patient’s global CNV distribution with the leftmost annotation bars indicating clinical subtypes (including gender) and the rightmost bar chart summarising CNV counts per patient (Fig. 4a). In total, we detected an average of 162 CNVs per individual, with deletions significantly outnumbering duplications (19,955 vs. 4,098; *p* < 2.2e-16). Copy number states reveal a dominance of deletions (Fig. 4b), whereas most CNVs ranged from 1 kb to 50 kb (Fig. 4c). Chromosome-wide analysis shows a general correlation between chromosome length and CNV burden, with chromosome 3 has the largest CNV count after normalizing with the length of chromosomes (Fig. 4d and Supplementary Fig. 14a). We also observed a good correlation between CNV count and number of genes in chromosomes (Pearson r = 0.56), in which chr1, chr4, chr19 and chr13 deviate from the trend (Supplementary Fig. 14b). Further clinical stratification indicates that neurofibromatosis type 1 (NF1) cases harbour the greatest CNV load, especially copy number loss (Fig. 4e). In addition, NF1 and epilepsy cases presented highest number of pathogenic and likely pathogenic CNVs, in consistent with their clinical severity (Supplementary Fig. 14c). Gender comparisons underscore a higher count of deletions and benign CNVs in male patients relative to females (Fig. 4f, g), but no difference is observed regarding to pathogenic CNVs (Supplementary Fig. 14d). Interestingly, the significant difference between females and males is only observed in DD patients, indicating the gender-biased CNV burden in DD (Fig. 4h).

**Fig. 4.**
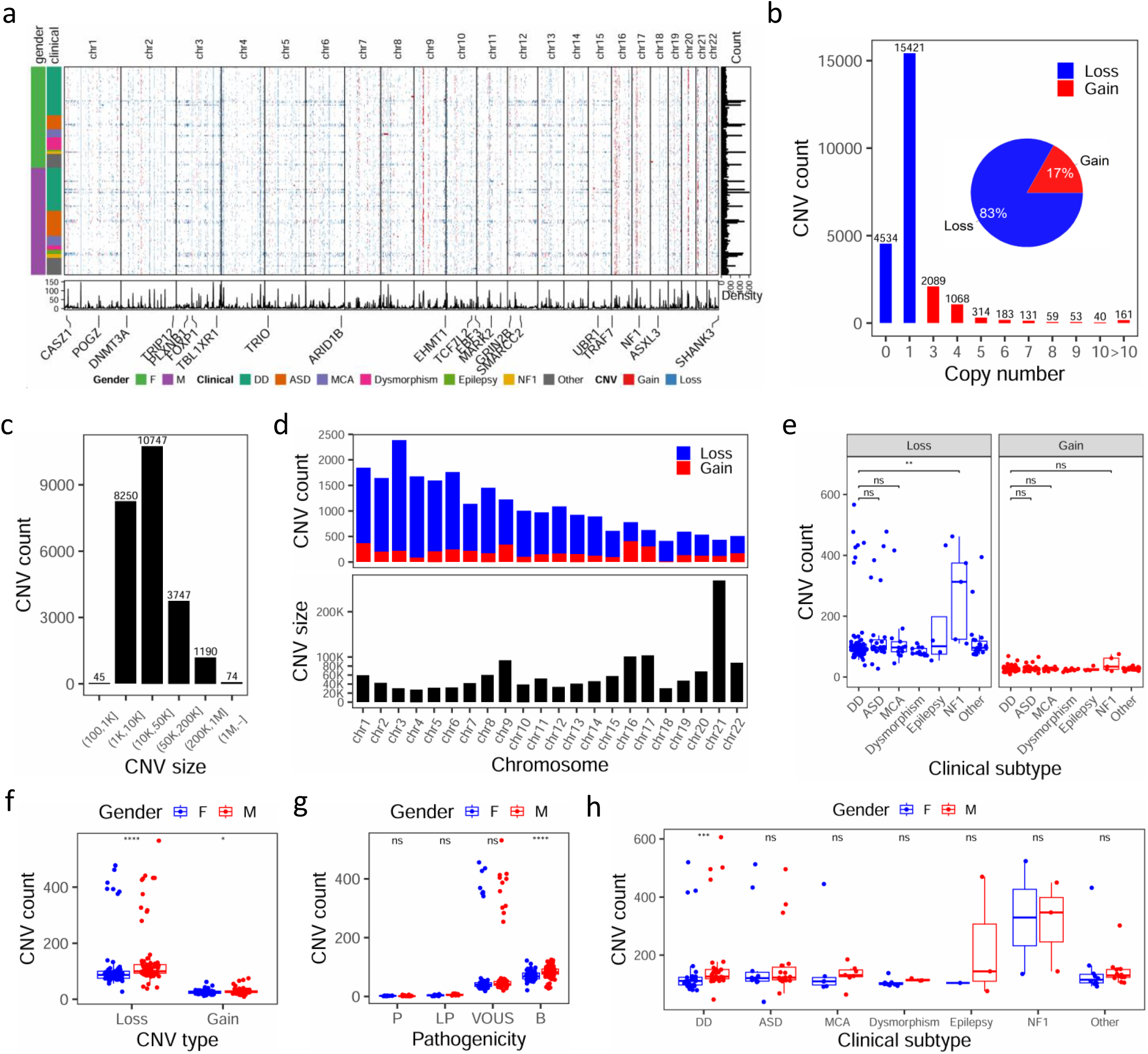
Genomic architecture of 150 paediatric patients with developmental defects analysed with CNVPipe. **a** Heatmap showing the distribution of CNVs identified from 150 patients. Columns represent genomic coordinates and rows represent patient samples. Leftmost bars indicate the phenotype of patients including their gender and clinical subtypes. Central bar plots indicate the number of CNVs detected from each patient. The rightmost bar indicates number of CNVs in each sample. Bottom density plot indicates the density of CNVs at different genomic locations. DD: developmental delay; ASD: autism spectrum disorder; MCA: multiple congenital anomalies; NF1: neurofibromatosis type 1. **b** Bar plot showing the distribution of CNVs with different copy numbers, pie plot showing the percentage of copy number loss and gain. **c** Bar plot showing the number of CNVs within different size ranges. **d** Bar plot showing the number and size of CNVs in each chromosome. **e** Box plots showing comparisons of CNV counts among patients with different clinical subtypes (see b for the full terms of the abbreviations used) divided by CNV type (gain and loss). **f** Comparison of CNV count between male and female patients grouped by CNV type. **g** Comparison of CNV count between male and female patients grouped by predicted pathogenicity (P: pathogenic; LP: likely pathogenic; VOUS: variant of unknown significance, and B: benign). **h** Comparison of CNV count between male and female patients grouped by clinical subtypes. Wilcoxon test *: *p* < 0.05, **: *p* < 0.01, ***: *p* < 0.001, ****: *p* < 0.0001, ns: not significant.

Among the clinical categories analysed, ASD showed the most striking CNV landscape^31^. Five individuals (IDs 27859, 28448, 28941, 29405, 29888) displayed an exceptionally high CNV burden relative to the remaining ASD cohort (Fig. 5a). These “high-burden” cases accounted for 36 of the 37 pathogenic or likely pathogenic events detected across all ASD genomes (36/37, *p* = 2.435e-7), consistent with the view that early, deleterious CNVs can trigger genome-wide instability and amplify mutation load (Fig. 5b). Functionally, 16 of the CNVs intersected well-established neuro-developmental genes, including *CASZ1, POGZ, DNMT3A, PLXNB1, FOXP1, TBL1XR1, TRIO, ARID1B, EHMT1, TCF7L2, EBF3, GRIN2B, SMARCC2, NF1, ASXL3,* and *SHANK3*, supporting direct genotype-phenotype links (Fig. 5c and Supplementary Table 3). In addition, CNVPipe uncovered eight intergenic, recurrent CNVs shared by ≥3 ASD patients but absent from population controls (Supplementary Table 4). Although annotated as benign or of uncertain significance, none overlapped coding sequences; overlay with UCSC transcription-factor binding sites and ENCODE cCREs revealed that these small intervals disrupt regulatory motifs for immunomodulatory factors such as *GATA3* (chr10) and *CEBPB* (chr20) (Fig. 5d and Supplementary Tables 5, 6). These observations point to an immune-regulatory axis in ASD pathogenesis that may be invisible to exon-centric analyses.

**Fig. 5.**
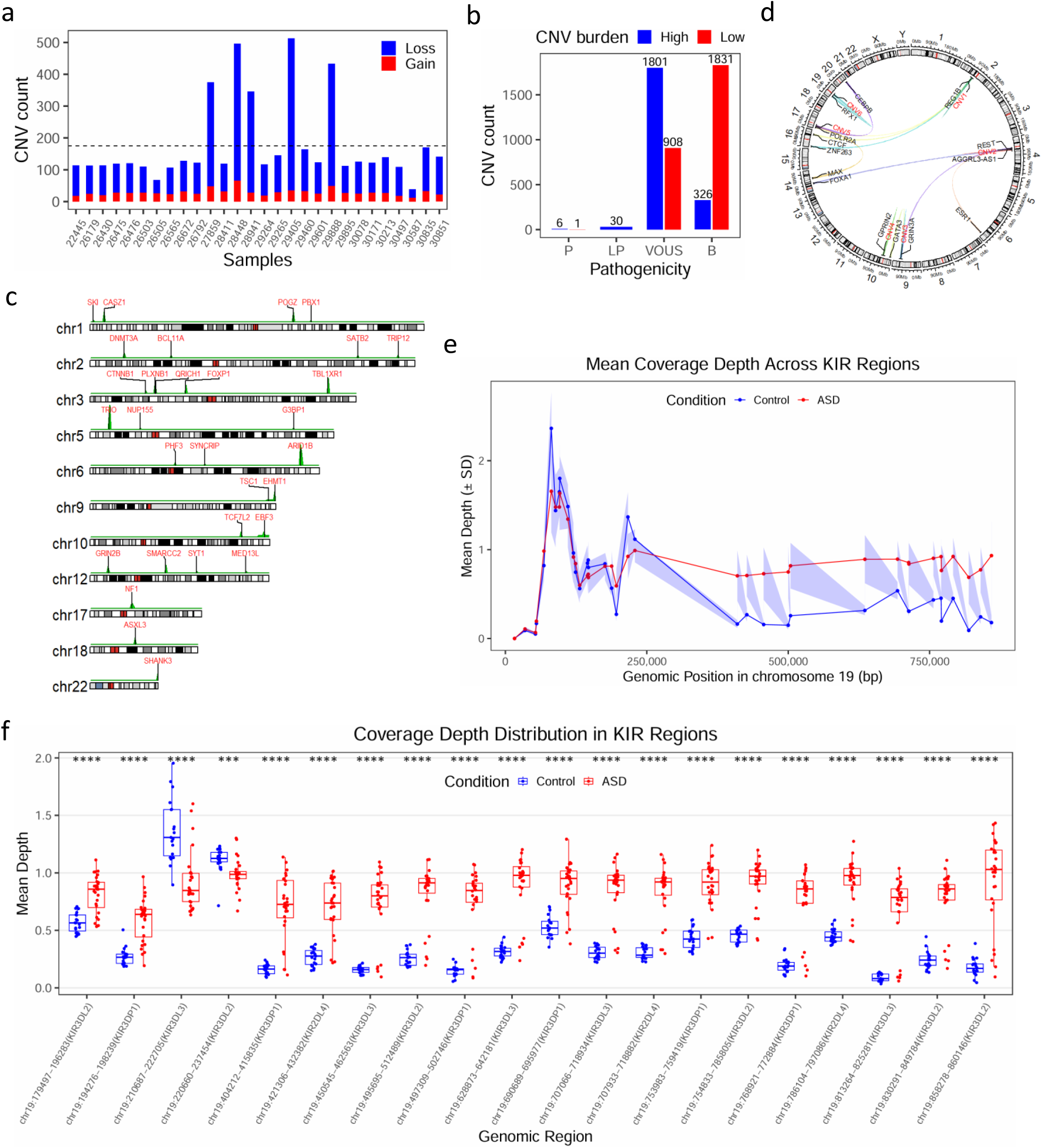
CNV detection in different genders and in patients with autism spectrum disorders (ASD). **a** Bar plot showing the CNV count in 28 ASD patients. **b** Bar plot showing the count of CNVs with different pathogenicity in high and low CNV burden and groups. **c** Karyoplot showing the genome-wide density of CNVs in high and low CNV burden and groups. **d** Karyoplot showing the CNVs that overlapped with reported ASD-risk genes. **e** Mean coverage depth across KIR regions in chromosome 19. **f** Comparison of coverage depth among ASD and control samples in 20 KIR regions. ***: *p* < 0.001, ****: *p* < 0.0001.

To probe this axis further, we examined copy-number architecture within the highly polymorphic human leukocyte antigen (HLA) and killer immunoglobulin-like receptor (KIR) loci, regions often poorly resolved by short-read pipelines. The depth-aware merging strategy by CNVPipe produced uniform coverage profiles across classical HLA genes, indicating no systematic dosage shift (Supplementary Fig. 15). In contrast, the KIR cluster on chr19q13.4 displayed pronounced copy-number variation in ASD genomes (mean coverage fold-change > 2 in high-burden cases; Fig. 5e, f and Supplementary Fig. 15). Because KIR genes modulate natural-killer-cell signalling, their dosage imbalance offers a tangible mechanistic link between innate immunity and neurodevelopment. Together, these results demonstrate how CNVPipe resolves complex immune-gene CNVs and integrates regulatory annotations, thereby delivering a functional-genomic framework for dissecting ASD and other complex disorders.

### Tracing tumour evolution through CNV score calculation from single-cell DNA sequencing (scDNAseq)

We employed Varbin^32^ method in the single-cell CNV-calling part of CNVPipe (CNVPipe-scDNAseq). Knowing the exact ploidies of single cells is critical to obtain accurate integer copy-number profile at the final step. However, most of the single-cell datasets lack of the results of fluorescence-activate cell sorting (FACS). Therefore, we propose a new way that utilises least-square rounding and peak-approaching strategy to accurately estimate ploidies for single cells (Supplementary Fig. 1). We evaluated our approach on a breast cancer dataset of 100 single-cells, whose ploidies are captured to be 1.7/2/3/3.3 by FACS^32^ (Supplementary Fig. 16). In this case, assuming all the cells to be diploid is apparently wrong, even though it can accurately cluster cells into four sub-populations (Fig. 6a). This can be misleading when prior ploidy information is absent. Least-square rounding method can accurately infer the existence of hapdiploid and aneuploid cells, at the expense of obtaining cells with highly variable ploidies (5, 6 or larger), leading to an extra sub-population as shown in the tail of density plot and UMAP plot (Fig. 6b). Our strategy approaches the cells into their most adjacent peak ploidies, which can provide precise inference of ploidies for all cells, eliminating the emergence of outliers (Fig. 6c). We utilised CNVPipe-scDNAseq workflow to analyse this dataset, and identified three clusters based on the CNV score calculated from integer copy number profile, including ‘near-normal’ cells, ‘cnv-low’ cells and ‘cnv-high’ cells (Fig. 6d). A phylogenetic tree was inferred by minimum spanning tree (MST) method using minimal event distance calculated from pairwise single cells (Fig. 6e). This tree accurately clustered cells into sub-population, and presented a branching evolutionary pattern.

**Fig. 6.**
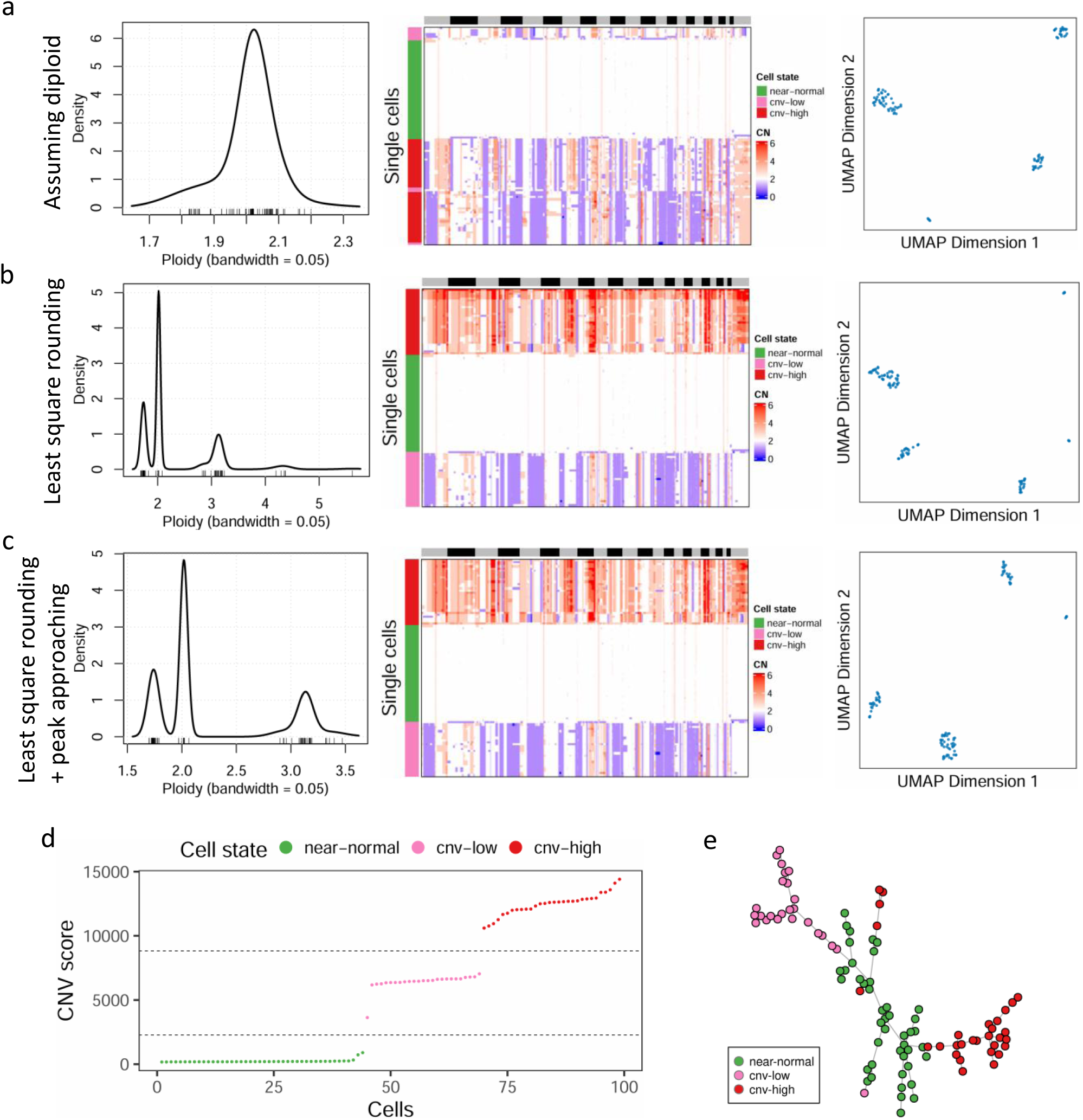
Evaluation of ploidy estimating approaches in cancer single cells by CNVPipe-scDNAseq and construction of cancer phylogenetic trees. Density plot showing the ploidies calculated from final integer copy-number matrix, heatmap showing the whole-genome distribution of CNVs in all single cells and UMAP plot showing the clustering of single cells based on integer copy number profile, when **a** all cells are assumed to be diploid; **b** using least-square rounding method to infer ploidies; and **c** using least-square rounding method in combination of peak-approaching strategy to infer ploidies. **d** One dimensional clustering separating the single cells into ‘near-normal’, ‘cnv_low’ and ‘cnv_high’ groups based on CNV score. **e** A phylogenetic tree with 100 single cells from a breast cancer tissue. Nodes represent cells, while green colour indicates near-normal cells, pink indicates cells with low number of CNVs, and red indicates cells with high number of CNVs.

To further explore the utility of CNVPipe-scDNAseq for clinical datasets, we analysed scDNAseq data from six triple-negative breast cancer patients who underwent neoadjuvant therapy^33^. Approximately 90 single cells were isolated before treatment, during treatment, and post-treatment (for one patient), and sequenced via single-nucleus sequencing. By applying our CNVPipe-scDNAseq workflow and MST-based phylogenetic tree construction, we found that in patients 1–3, cells occupying the final branches of the tumour phylogeny (red ellipse, in Fig. 7a-c) originated exclusively from pre-treatment samples, suggesting the tumour was effectively eradicated by therapy. In contrast, the end-phase branches in patients 4–6 contained cells from pre-, mid-, and post-treatment stages (blue ellipse, in Fig. 7d-f), implying surviving cancer cells and a poor clinical response. These findings align with patient outcomes and underscore the precision of our pipeline in evaluating therapeutic efficacy.

**Fig. 7.**
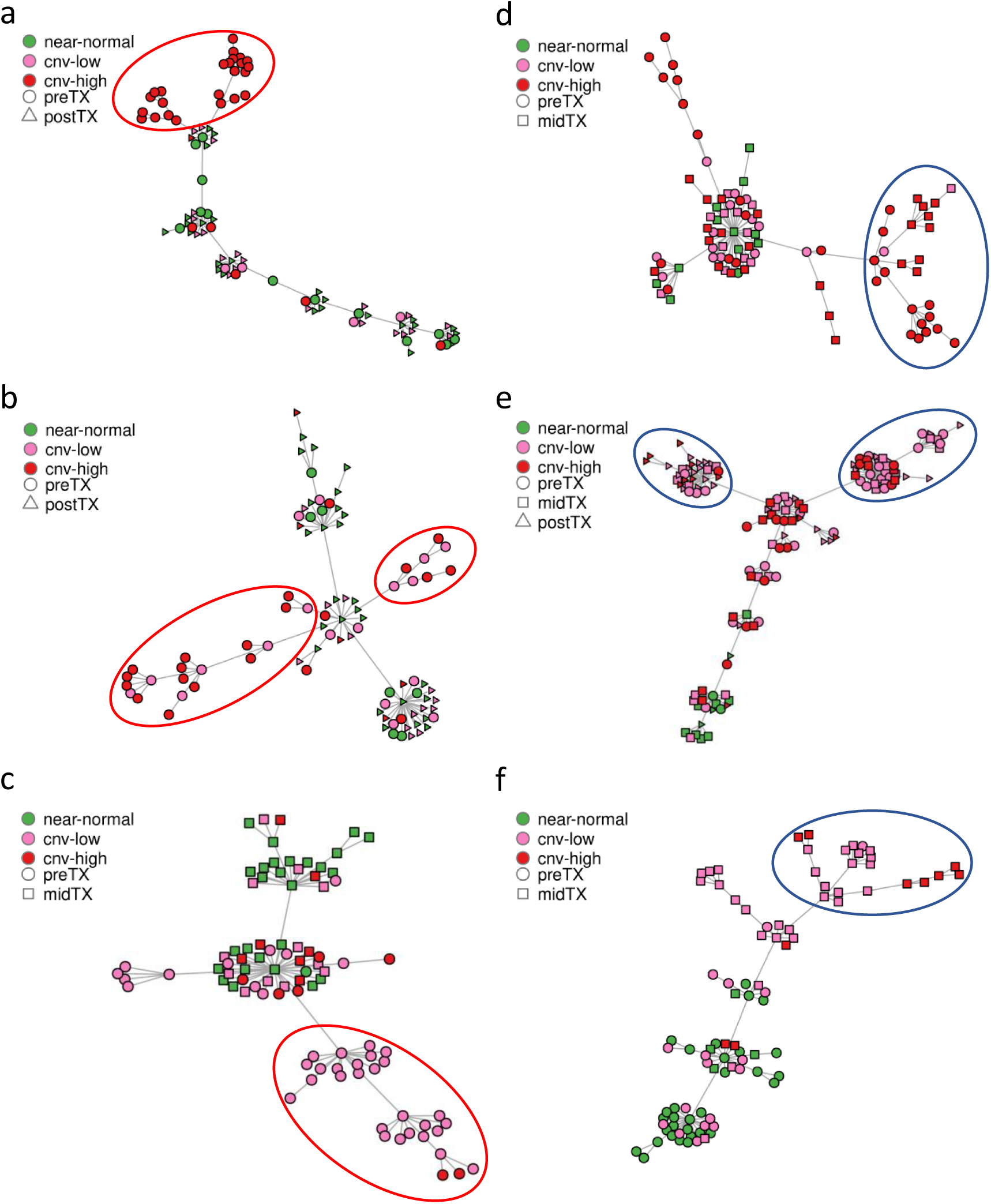
Phylogenetic trees of six datasets from triple-negative breast cancer patients who underwent neoadjuvant therapy were analysed by CNVPipe-scDNAseq. Six samples have **a** 93, **b** 90, **c** 92, **d** 93 **e** 138 and **f** 91 cells, respectively. Nodes in the tree represent cells, colour represents cell groups based on CNV score, circle represents cell from sample before treatment (preTX), square represents in the middle or post treatment (midTX or postTX), triangle represents post-treatment (postTX).

Overall, these results highlight the adaptability of CNVPipe across a spectrum of data types (Fig. 8) from low-depth single-cell assays and clinical WGS to high-depth WGS in stem cell models, underscoring its potential in elucidating disease mechanisms, advancing developmental and gene-editing research, and enhancing our understanding of cancer evolution.

**Fig. 8.**
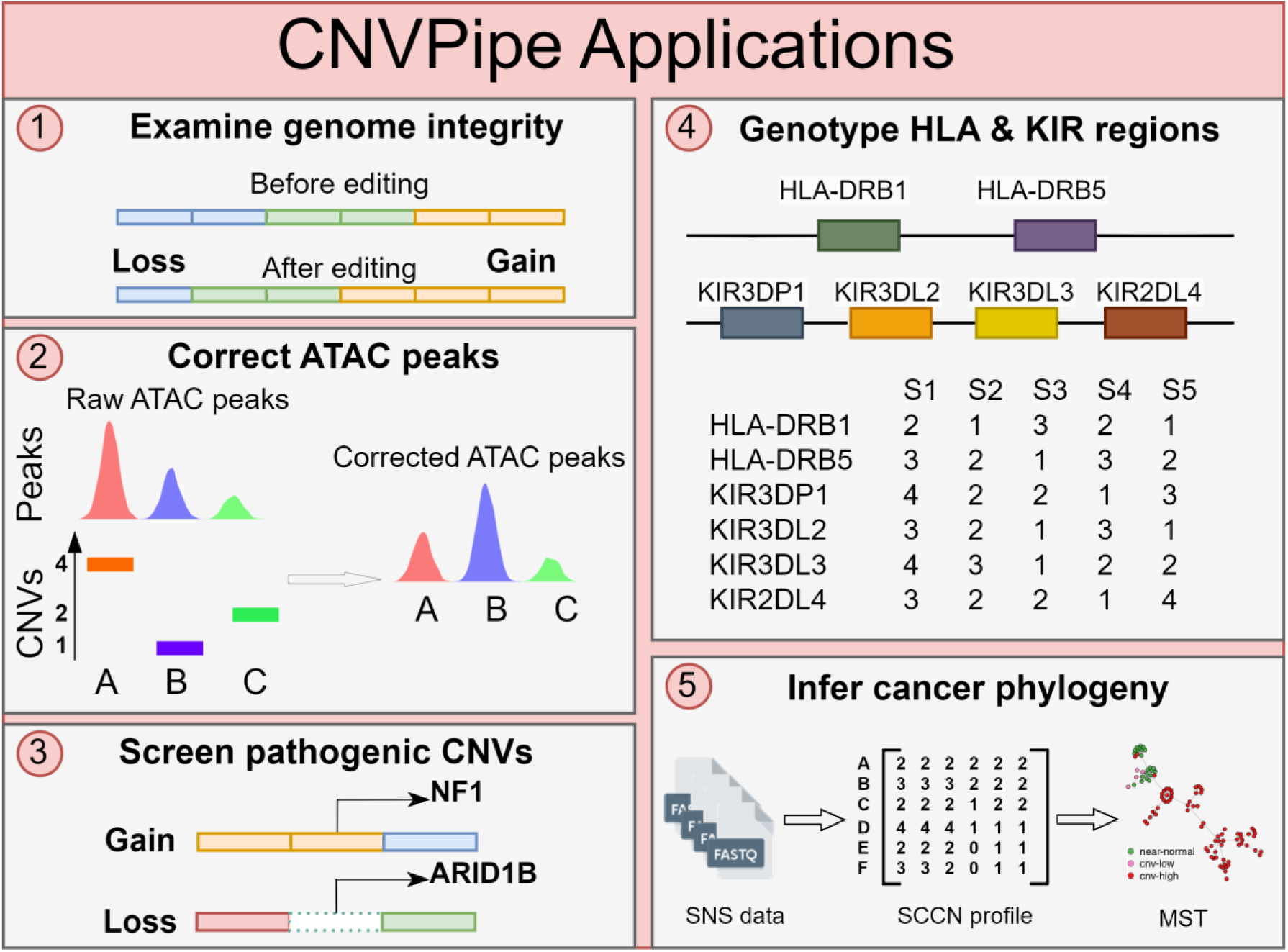
Schematic diagram presenting the broad applications of CNVPipe. CNVPipe is applicable for examining the genome integrity after gene editing, correcting for differential accessible ATAC peaks, screening pathogenic CNVs from clinical samples, genotyping complex HLA and KIR regions and assisting for the inference of cancer phylogeny.

## Discussion

In this study, we present CNVPipe as a streamlined solution for CNV detection, aiming to address the persistent challenges of high FDR and diverse sequencing depths in WGS data. By integrating multiple CNV-calling tools and implementing a machine learning-based classifier, CNVPipe demonstrates solid performance across a range of CNV sizes and sequencing depths. Notably, the Snakemake-based architecture of CNVPipe provides a user-friendly workflow that consolidates raw data pre-processing, CNV detection, downstream annotation, and automated visualization.

One of the core strengths of CNVPipe is its adaptability. Ensemble-based merging strategies enable high accuracy under conditions where individual tools might otherwise fail— such as at low sequencing depths with large CNVs, or high depths with subtle, smaller events. Our results indicate that combining read-depth, split-read, and read-pair methods can yield more balanced recall-precision trade-offs than relying on individual algorithms alone. Furthermore, the use of simulated data and empirical WGS benchmarks reveals that CNVPipe, particularly its SVM variant (CNVPipe-SVM), maintains robust detection while minimizing spurious calls. Although some read-depth-based approaches underperform for certain real-world datasets, we attribute this to biases in “ground truth” structural variant sets, which often favour smaller deletions over more complex or larger-scale variations. A broader, more representative reference set could enhance training and improve performance for both read-depth-based callers and classifiers of CNVPipe.

Beyond short-read WGS, our pipeline is poised for future enhancements. The modular Snakemake backbone of CNVPipe allows integrating additional CNV and SV tools, making it suitable for specialized tasks such as exome-based CNV detection, long-read sequencing analysis, or more comprehensive SV discovery. We also plan to refine its visualisation capabilities by introducing an interactive genome viewer that can highlight genomic loci, previously reported CNVs, and relevant annotations, thereby streamlining user validation.

Applying CNVPipe to paediatric patients with developmental defects illustrates its capacity to detect recurrent CNVs that could underlie disease phenotypes. The ability of our pipeline to identify and preliminarily validate these regions through manual inspection offers a foundation for follow-up experiments (e.g., Sanger sequencing or droplet digital PCR) to confirm breakpoints and copy number states. Such in-depth studies could shed new light on the contribution of specific CNVs to neurodevelopmental anomalies and pinpoint novel genetic targets for clinical testing.

We also explored the power of CNVPipe for single-cell genomics, deploying an MST-based method to produce phylogenetic trees that track tumour evolution. Our analyses will further correlate the resulting phylogenies with clinical outcomes and highlight how single-cell datasets can illuminate evolutionary processes, therapy resistance, and heterogeneity within tumours.

Looking ahead, incorporating long-read sequencing data will likely expand the ability of CNVPipe to resolve complex or repetitive genomic regions. In parallel, deep learning approaches could improve variant calling by learning nuanced features across larger training sets. Further opportunities exist in single-cell RNA-sequencing data, where methods such as inferCNV^34^, Sitka^35^, Numbat^36^, or CopyKAT^37^ could complement or integrate with CNVPipe, potentially enabling multi-omic perspectives on tumour or developmental processes.

Overall, our work underscores the flexibility of CNVPipe as a comprehensive platform for CNV detection across multiple sample types and clinical applications. By offering robust ensemble calling, machine learning-assisted filtering, and specialized single-cell modules, CNVPipe has the potential to advance genomics research and precision medicine in fields ranging from developmental disorders and stem cell models to cancer biology.

## Methods

### Design of CNVPipe

CNVPipe is a Snakemake-based pipeline that integrates five popular CNV-calling tools, each using different CNV signals and statistical methods. CNVPipe computes multiple scoring metrics to refine and prioritise high-confidence CNVs, and integrates a pathogenicity prediction method based on 2019 ACMG standards^38^. For population-level studies, CNVPipe enables the identification of recurrent CNVs among individuals with specific phenotypes. The pipeline generates a user-friendly report that includes finalised CNV tables and plots displaying the distribution of reads and BAF information for manual confirmation. The minimum input for CNVPipe is a configuration file and fastq files of the original sequencing reads. Figure 1 provides a step-by-step workflow of CNVPipe. This design ensures stability across different machines and environments, making CNVPipe a valuable tool in the CNV study field.

### Calling and merging of CNVs

- **Pre-processing**: CNVPipe, our pipeline for CNV calling, performs pre-processing using fastp^39^ to remove adaptors and filter out reads with poor quality. It then uses bwa^40^ to map reads to the human reference genome and generate bam files. GATK4^41^ is used to recalibrate base quality and mark duplicate reads, producing bam files that are ready for CNV calling.
- **Determining bin size (optimal)**: Determining an appropriate bin size and resolution is crucial for accurate and sensitive CNV calling. Bin size can significantly influence the analysis, and it is important to choose the right resolution for data with different sequencing depths. Our simulation studies have recommended resolutions for various sequencing depths (Supplementary Table 8). Another option is to use CNVPipe, which utilises a weighted median method to estimate the resolution for the user’s data. First, CNVPipe identifies the sample with the median number of reads among all samples. Then, it calculates the read depth in each chromosome and calculates a weighted median read depth across the genome. Finally, CNVPipe determines the resolution by ensuring that each bin is covered by 15,000 nucleotide bases, which is equivalent to approximately 100 reads for paired-end 150-bp sequencing data.
- **Parallel CNV calling**: CNVPipe integrates five widely cited CNV-calling tools for parallel CNV detection. These include three read-depth-based methods – CNVKit^8^, CNVpytor^9^ and cn.MOPS^6^ – as well as two approaches that utilize split reads and pair-end information – Delly^14^ and Smoove^13^. All tools are actively maintained and have been widely adopted in recent studies. The read-depth-based tools employ distinct normalization and segmentation strategies. For example, CNVKit corrects for GC content and repetitive regions using a rolling median approach. CNVpytor applies a mean-shift algorithm for bin segmentation and utilizes t-tests to identify high-confidence CNV regions. cn.MOPS models read depth across samples using a mixture of Poisson distributions for each bin, segments bins based on informative versus non-informative calls, and infers CNVs both across samples and along chromosomes. Smoove streamlines and accelerates CNV calling for short-read data by integrating Lumpy and other utilities, while Delly employs a two-step approach that sequentially analyzes discordant paired-end reads and split reads to detect CNVs.
- **SNP calling**: For SNP calling, CNVPipe integrates GATK4 and Freebayes^42^, which will be used for refining CNVs based on BAF. GATK4 HaplotypeCaller will be used to call SNPs if the depth of data is high (> 10×), while Freebayes will be used if the depth is low.
- **Merge CNVs from all tools**: Given that the performance of CNV-calling tools can vary substantially depending on sequencing depth, CNVPipe implements depth-dependent merging strategies to optimize integration of results. CNVs are merged recursively when their reciprocal overlap exceeds 0.75, with the merged CNV defined by the outermost breakpoints of the overlapping events. The copy number estimate for the merged CNV is derived from read-depth-based methods. To track the merging process, the overlap fraction and contributing tools are recorded. As the order of merging may influence the final outcome, the merging sequence is determined based on sequencing depth. For samples with a mean read depth below 1×, results from cn.MOPS, CNVKit, and CNVpytor are combined. For depths between 1× and 5×, the merging incorporates calls from cn.MOPS, CNVKit, Delly, CNVpytor, and Smoove. For data exceeding 5× depth, the merging prioritizes Smoove, Delly, CNVKit, CNVpytor, and cn.MOPS. This approach ensures that precise breakpoints from split-read methods and accurate copy number estimates from depth-based approaches are both preserved.

The parameters used in CNVPipe for CNV calling and SNP calling are described in Supplementary Methods.

### Scoring metrics

Scoring metrics play a critical role in refining and prioritising high-confidence CNVs during and after the merging process. CNVPipe utilizes the following scoring strategies to assess CNV reliability:

- Accumulative score (AS): This metric quantifies the cumulative overlap fractions of CNVs detected by multiple tools during merging. A higher AS indicates that a CNV was consistently identified by more tools, suggesting greater confidence in its validity.
- Depth score (DS): CNVPipe computes DHFFC and DHBFC using duphold (https://github.com/brentp/duphold) to evaluate depth fold-changes relative to flanking regions (DHFFC) and to GC-matched bins (DHBFC). Based on these values, CNVPipe assigns a DS for each CNV, adjusted according to CNV type (deletion or duplication). Higher DS scores correlate with increased confidence in CNV detection.
- Allele frequency score (AF): CNVPipe leverages BAF (B Allele frequency) information via CNVfilteR^43^ to prioritize CNVs supported by SNP data. Each CNV is assigned an AF score reflecting its concordance with SNP-based evidence. This method depends on accurate SNP calling and requires sufficient sequencing depth; in cases of insufficient BAF data, CNVfilteR may default to labeling CNVs as ‘True’.
- Good-region score (GS): The human genome contains numerous low-complexity and highly repetitive (LCHR) regions that are prone to false-positive CNV calls. CNVPipe evaluates the overlap between merged CNVs and a custom list of LCHR regions, assigning a GS to each CNV. A higher GS indicates reduced overlap with such problematic regions, implying greater CNV reliability.
- Mappability score (MS): CNVPipe finds overlap between merged CNVs and a custom list of low-mappability genomic regions, and assigns a MS to each CNV. A higher MS implies less overlap with low-mappability regions, indicating more reliable CNVs.
- Normal score (NS): CNVPipe integrates high-frequency CNV data from dbVar (NCBI) to identify common variants in the general population. It calculates the overlap between merged CNVs and these benign CNVs, generating an NS based on cumulative overlap fractions. A higher NS suggests lower similarity to common, likely non-pathogenic, CNVs, potentially indicating pathogenic relevance. Although NS is not primarily used for filtering low-confidence calls, it can aid in interpreting CNV pathogenicity.

A detailed description of these scoring metrics is provided in the Supplementary Methods.

### Predicting pathogenicity

To predict the pathogenicity of merged CNVs, CNVPipe incorporates ClassifyCNV^44^, a tool that applies the CNV classification guidelines established by the American College of Medical Genetics and Genomics (ACMG) and integrates multiple genomic databases to evaluate pathogenicity. ClassifyCNV categorizes CNVs into one of five classes: pathogenic, likely pathogenic, benign, likely benign, or uncertain significance. Following pathogenicity prediction, CNVPipe assigns each CNV a “pathogenicity score” (PS), which is used to prioritize variants with higher likelihoods of clinical relevance.

### Finding recurrent CNVs

To identify recurrent CNVs in population-level studies, CNVPipe implements an iterative merging strategy. Initially, all CNVs detected across samples are collected and subjected to recursive pairwise comparison. If two CNVs exhibit a reciprocal overlap greater than 0.8, they are merged into a new CNV defined by the innermost breakpoints of the original events, and the two original CNVs are removed from the list. If the overlap threshold is not met, the original CNVs are retained, and the merging process continues iteratively until all CNVs have been evaluated. During this process, the number of samples supporting each merged CNV is recorded, enabling users to filter recurrent CNVs based on their specified frequency thresholds. This strategy facilitates the association of recurrent CNVs with specific phenotypes in large-scale cohort analyses.

### Data simulation

To evaluate the performance of CNVPipe and other CNV detection tools, we developed a custom CNV simulator (https://github.com/sunjh22/CNV-simulator), which generates synthetic whole-genome datasets with known CNVs serving as a ground-truth reference. Users can specify the desired number and size distribution of CNVs, with CNV lengths modeled according to an exponential distribution with parameter λ. The simulator randomly assigns copy number alterations to paternal and maternal chromosomes and modifies the reference genome accordingly, enabling haplotype-specific CNV simulation. This approach allows for the generation of complex events such as heterozygous deletions or amplifications, resulting in total copy numbers of 0, 1, 3, 5, and so on. Whole-genome sequencing (WGS) reads are then simulated using ART_illumina^45^. To avoid CNV placement in low-complexity or highly repetitive (LCHR) regions, we provided a list of genomic accessible regions as input to the simulator.

We generated datasets across multiple CNV size levels (λ = 10 kb, 50 kb, 200 kb, and 1 Mb) to assess the impact of CNV size on detection accuracy. For each size category, six sequencing depth levels (0.1×, 0.5×, 1×, 5×, 10×, and 30×) were evaluated, with six replicate samples per group, to systematically investigate the influence of sequencing depth on CNV calling performance.

To evaluate CNVPipe’s ability to identify recurrent CNVs, we compiled a set of 50 pathogenic deletions and 50 duplications larger than 10 kb from the dbVar pathogenic CNV database. We then simulated 30 samples, each harboring these 100 recurrent CNVs, along with an additional 100 sample-specific CNVs. Sequencing reads were generated at three depth levels (1×, 10×, and 30×) for downstream analysis.

### Collecting WGS data for benchmark

We obtained eight real WGS datasets including NA12878-1, NA12878-2, CHM13, AK1, HG002, HG00514, HG00733, and NA19240 for benchmarking purposes. To ensure a high level of confidence in the CNV set used as ground truth, we selected samples with high-quality CNV annotations. These samples include NA12878 (part of the 1000 Genomes Project), CHM13 (a haploid human genome), AK1 (a de novo assembled Korean genome), HG002 (from the Genome in a Bottle (GIAB) project), and three other samples from the Human Genome Structural Variation Consortium. The original fastq data and relative CNV truth sets were downloaded from the supplementary file of the original paper or from the NCBI dbVar database. Only high-confidence CNV sets were downloaded for all samples, and we selected only deletions and duplications with sizes over 1 kb to generate a ground-truth set for CNVs.

Detailed information about real WGS dataset can be found in Supplementary Table 9.

### Performance evaluation

We applied two criteria to define true positive (TP) CNVs for performance evaluation. In the first method, an identified CNV was classified as a TP if it exhibited greater than 0.8 reciprocal overlap with a ground-truth CNV and shared the same CNV type (e.g., deletion or duplication). This criterion was specifically designed for benchmarking on real whole-genome sequencing (WGS) data, where ground-truth CNVs are derived from structural variant (SV) datasets that typically provide only CNV type information without exact copy number values. The second method imposed a stricter requirement: in addition to reciprocal overlap > 0.8 and concordant CNV type, the predicted CNV also had to match the exact copy number value of the ground-truth CNV. Simulation results were evaluated using the second method, while empirical results were assessed by the first method.

Three ensemble CNV-calling tools – sv-callers, ClinSV and MetaSV, and five individual CNV callers incorporated in CNVPipe were evaluated with default parameters. To comprehensively evaluate the performance of CNV-calling methods, we employed multiple metrics. Accuracy was defined as the proportion of correctly identified CNVs relative to all ground-truth CNVs. False discovery rate (FDR) represented the proportion of false positives among all called CNVs. The F1 score, calculated as the harmonic mean of accuracy and precision (where precision = 1 – FDR), provided a balanced measure reflecting both sensitivity and specificity.

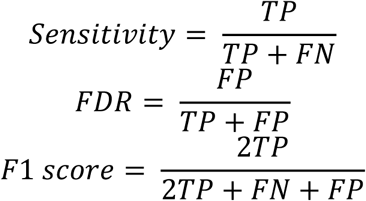

### Machine learning classifiers in CNVPipe

To enhance CNV filtering and prioritization, CNVPipe incorporates a suite of machine learning (ML) classifiers for distinguishing true positive (TP) CNVs from false positives (FPs). The implemented models include logistic regression, linear support vector machine (SVM), radial basis function (RBF)-kernel SVM, Gaussian process classifier, random forest, neural network, and Naïve Bayes. Training data were generated by comparing CNVPipe’s CNV calls against ground-truth sets derived from both simulated and real whole-genome sequencing (WGS) datasets. Each CNV was labeled as TP or FP and represented using a feature vector comprising the following variables: CNV size (size), copy number (CN), GC content (GC), contributing tools (tools), number of tools (ToolNum), accumulative score (AS), depth score (DS), allele frequency score (AF), good-region score (GS), mappability score (MS), DHFC, DHBFC, and DHFFC.

For simulated data, results from five samples were used for training, with the remaining sample reserved for testing. For real WGS data, seven randomly selected samples formed the training set, while the remaining sample served as the test set. The SVM model was trained using five-fold cross-validation and subsequently applied to predict TPs among CNVPipe’s output. Model performance was evaluated in terms of accuracy and false discovery rate (FDR), calculated by comparing predicted TPs with the ground-truth CNVs.

### DNA preparation of patient samples

Peripheral blood samples were collected from 150 paediatric patients (77 male, 73 female) with different types of developmental defects: 66 with developmental delay (DD), 28 with autism spectrum disorder (ASD), 13 with multiple congenital anomalies (MCA), 12 with dysmorphism, 4 with epilepsy and 27 with other phenotypes. In addition, blood samples were collected from 19 healthy donors to serve as reference controls. Genomic DNA was isolated using QIAamp® genomic DNA kits (Qiagen) and quantified by Qubit 2.0 fluorometer (Life Technologies). The integrity of the genomic DNA was assessed with 0.7% agarose gel electrophoresis.

### Library preparation and whole-genome sequencing

The sequencing libraries was prepared using the MGIEasy PCR-Free Library Preparation Kit following the manufacturer’s instructions. Briefly, 1 μg DNA sample was used as the starting material, followed by fragmentation with specific enzymes, and selection of the resulting product with size around 400-600 bp using magnetic beads. Then, adaptors were inserted into the fragmented DNA through end repair, A-tailing, and adaptor ligation. Adaptor-ligated DNA was amplified for eight cycles by PCR. The PCR product was purified by magnetic beads, and subjected to heat-denaturation along with a special molecule that is reverse-complementary to one of the PCR product strand. The single-strand molecule was ligated using a DNA ligase to form circularized product. The circularized DNA was further digested by Exo enzyme and subjected to a further purification. The final DNA concentration was measured using Qubit ssDNA Assay.

Sequencing was conducted according to the MGISEQ-2000RS protocol provided by the manufacturer. To construct DNA NanoBalls (DNB), the single-strand circular DNA library was PCR-amplified and quantified using a Qubit fluorometer with an expected minimum concentration of 10 ng/μl. The constructed DNBs were loaded onto the sequencing chip and the sequencing procedure was initiated after installing the sequencing chip with preloaded reagents. All samples were sequenced in PE150 mode (300 cycles of run), and the minimum physical coverage equivalent to 10-fold (10x) base coverage was set to meet the requirement for the detection of copy number variations and minimize false-negative results. The binary file, which contains nucleotide bases and quality scores, was converted into FASTQ format with Phred+33 quality score, pending for downstream bioinformatics analysis.

### Manual inspection of CNVs

CNView were used to manually check all the CNVs identified in iPSC P5, iPSC P30, dLNC93 and control samples by both CNVPipe-SVM and sv-callers. CNVs with q value less than 10^-4 were determined to be true, otherwise false. Samplot^46^ was also used to re-confirm these CNVs. Recurrent CNVs identified from paediatric patients and CNVs overlapped with regulatory elements were also manually validated using the aforementioned approach.

### iPSC culture and RGC differentiation

Human iPSCs (ALSTEM) are cultured in growth medium consisting of mTeSR1 medium with completed supplements (STEMCELL Technologies). Once cells reached 80% confluence, they were subcultured into new culture plates pre-coated with Geltrex (Gibco), which was diluted in DMEM/F12 (Gibco) at a 1:50 ratio.

RGC differentiation started from embryoid body (EB) formation by dissociating undifferentiated iPSC colonies with ReLeSR (STEMCELL Technologies) and suspending the small cell aggregates in ultra-low attachment 6-well plates with DMEM/F-12 medium containing 20% knockout serum replacement (KSR), 1X GlutaMAX, 1X nonessential amino acids (NEAA) and 0.1 mM β-mercaptoethanol for 1 week. Then, EBs were transferred to gelatin-pretreated plates and cultured in DMEM/F-12 medium containing 20% KSR, 1X GlutaMAX, 1X NEAA, 0.1 mM β-mercaptoethanol, 10% fetal bovine serum (FBS) and 10 ng/ml basic fibroblast growth factor (bFGF) for 1 week. Neural rosettes would appear during this period. The neural rosettes were mechanically lifted with a syringe needle and a pipette tip and were transferred to ultra-low attachment 6-well plates, and grown in suspension in RGC differentiation medium (DMEM/F-12 medium containing 20% KSR, 1X GlutaMAX, 1X NEAA, 0.1 mM β-mercaptoethanol, 10% FBS, 10 ng/ml bFGF and 10 μM DAPT) for 5 days to allow the formation of neurospheres. Next, the neurospheres were transferred to laminin-coated plates with RGC differentiation medium containing fresh DAPT (STEMCELL Technologies) that was provided every other day. On day 40, the RGCs were harvested by accutase treatment for subsequent investigation.

### CRISPR-Cas9-mediated deletion of target gene

A long non-coding RNA named LNC000093 was selected as a target gene for knockout in iPSCs. The Alt-R CRISPR-Cas9 system (IDT) was used to generate LNC000093-deletion in iPSCs. A pair of custom sgRNAs targeting two genomic regions flanking LNC000093 was designed with proprietary algorithms of IDT. The sgRNAs were mixed with Alt-R® S.p. Cas9 Nuclease V3 (IDT) to form ribonucleoprotein (RNP) complexes, which were subsequently used to assemble transfection complexes by incubation with Lipofectamine RNAiMAX reagent (Invitrogen). In total, 3 × 10^5^ cells per well in 24-well plates were transfected with 10 nM RNP complex. A pair of PCR primer was designed to target the regions flanking the deletion region of LNC000093. Genomic DNA was extracted from 1 x 10^6^ CRISPR-edited cells using PureLink Genomic DNA Mini Kit (Invitrogen) following the manufacturer’s instructions. 50 ng DNA was employed in each PCR reaction with the use of Platinum SpuerFi II Green PCR Master Mix (Invitrogen) according to following cycling conditions: 98°C for 30 s; 30 cycles of 98°C for 15 s, 60°C for 15 s, 72°C for 2.5 min; 72°C for 5 min. The PCR amplicons were then used in 2% agarose gel electrophoresis to assess the CRISPR-deletion effect based on the product size difference.

### RT-qPCR

Total RNA was isolated using a Trizol/RNeasy hybrid method which involved the use of TRIzol reagent (Life Technologies) and the RNeasy Mini Kit (QIAGEN). The RNA’s quantity and quality were assessed using a NanoDrop spectrophotometer.

ReverAid First Strand cDNA Synthesis Kit (ThermoFisher Scientific) was used to generate first-strand cDNAs, utilising 0.5-1 μg of total RNA per sample for all RT reactions. The resulting cDNA served as the input template for real-time qPCR, which was performed on a ViiA7 real-time cycler (Applied Biosystems) using the QuantiNova SYBR Green PCR mastermix (QIAGEN) according to the manufacturer’s instructions. In brief, the PCR reaction began with initial activation at 95°C for 2 min, followed by 40 cycles of 95°C for 30 s and 60 °C for 1 min. Melting curve analysis was incorporated in each run to monitor the specificity of reaction. Relative changes in gene expression between samples were calculated using 2^-ΔΔCt^ method, with GAPDH expression serving as the control for normalization.

### Droplet digital PCR

The genomic DNA of iPSC samples was extracted using PureLink Genomic DNA Mini Kit (Invitrogen) according to the manufacturer’s instructions. Droplet digital PCR (ddPCR) assay was performed using EvaGreen fluorescent nucleic acid dye to detect positive droplets. Each ddPCR reaction consisted of 10 μL 2X ddPCR EvaGreen Supermix (Bio-Rad), 250 nM forward and reverse primers, 20 ng gDNA sample, and nuclease-free water to top up the total reaction volume to 20 μL. The reaction mix was mixed thoroughly by short vortexing and was allowed to equilibrate at room temperature for 3 min before oil droplet generation. Then, 20 μL reaction mix and 70 μL Droplet Generation oil for EvaGreen (Bio-Rad) were loaded in a DG8 Cartridge (Bio-Rad), which was placed into the QX200 Droplet Generator (Bio-Rad) to produce droplets for PCR. 40 μL of droplet mix was then transferred to specified 96-well PCR plate (Bio-Rad). The plate was sealed with a heat seal foil using PX1 PCR plate Sealer (Bio-Rad). The thermal cycling reaction was conducted using Veriti Thermal Cycler (Applied Biosystems) under the following conditions: initial activation at 95°C for 5 min, followed by 40 cycles of 95°C for 30 s and 60°C for 1 min. Then, a signal stabilization step was carried out at 4°C for 5 min and 90°C for 5 min, with the reaction finally held at 4°C until data acquisition. A ramp rate of 2°C/s was applied throughout the cycling process to ensure the oil droplets reached the appropriate temperature. The ddPCR signals were read by the QX200 Droplet Reader and analyzed using the QuantaSoft Software (Bio-Rad).

### Construction of tumour phylogenetic trees

Single-cell DNA sequencing data of breast cancers were downloaded from NCBI SRA repository using accession number of SRP002535^32^ and SRP116771^33^. The raw fastq data was analyzed by CNVPipe-scDNAseq workflow to obtain integer copy number matrix. To calculate CNV score, bins with copy number 1 or 3 were given 1 point, copy number 0 or larger than 3 were given 2 points, while copy number 2 was given 0 point. CNV score was calculated by summing up the points in all bins. For each tumor sample, cells are sorted by CNV score in ascending order, and clustered into three groups by one-dimensional clustering method Ckmeans.1d.dp^47^. We then defined the group 1 as ‘near-normal’ cells, group 2 as ‘cnv-low tumor’ cells and group 3 as ‘cnv-high tumor’ cells. Unrooted undirected minimum spanning tree (MST) was constructed based on minimal event distance calculated from integer copy number profile of pairwise cells by using the implementation of Prim’s algorithm. Overall distance is guaranteed to be minimized and nodes are allowed to have more than two adjacent nodes in the tree. The time complexity of the algorithm is *O*(V^2^ + E), where V being the number of vertices (cells) and E being the number of edges. R package igraph^48^ (v1.3.2) was used to visualize the tree.

## Data availability

The simulated data used in this study were generated by the CNV-simulator software (see Methods section for details). The raw sequencing data for the real samples analysed in this study were downloaded from NCBI SRA or EBI ENA databases with the following accession IDs: NA12878-1 (SRR622457), NA12878-2 (ERR194147), CHM13 (SRR3986881), AK1 (SRR3602759), HG002(GIAB ftp site, ftp://ftp-trace.ncbi.nlm.nih.gov/giab/ftp/data/AshkenazimTrio/HG002_NA24385_son/), HG00514 (ERR894729, ERR894730, ERR899717, ERR899718, ERR903030), HG00733 (ERR899724, ERR899725, ERR899726, ERR903031), and NA19240 (ERR894723, ERR899710, ERR899711). As the single fastq file for HG00514, HG000733 and NA19240 cannot reach 30×, we downloaded several of them and concatenated. The raw scDNA-seq data were downloaded from NCBI SRA with accession IDs: SRP002535 and SRP116771. The raw sequencing data for 150 pediatric patients can be accessed in NCBI SRA with PRJNA1259217.

## Supplementary materials

Supplementary figures, tables, and methods can be found online.

## Acknowledgments

Studentship of J.S. was supported by a collaborative scheme with Southern University of Science and Technology (internal funding of The Hong Kong Polytechnic University). The clinical studies were supported by the Health and Medical Research Fund in Hong Kong (No. 16172331) and departmental funding schemes (Department of Health Technology and Informatics). Fellowship of J.S. and N.K.W. were funded by InnoHK initiative from the Government of the Hong Kong Special Administrative Region (Health@InnoHK Fund), and supported by the Centre for Eye and Vision Research (CEVR RP1.6) and The Hong Kong Polytechnic University (RiFood-CD55). W.J. was supported by National Natural Science Foundation of China [32170646, 32370688]. S.P.Y. was supported by 99QP and E-RD86.

## Author contributions

C.-L.H., S.P.Y. and W.J. conceived and supervised the project. J.S. developed and implemented the algorithms, and wrote the first draft of the manuscript. N.K.W. and Z.J. conducted the experiments and validations. H.-M.L. and F.M.L. coordinated the collection of patient samples and the demographic data. D.D., S.Z. and R.C.B.W. did the data analysis and biological interpretation. J.S. and C.-L.H. completed the figures and manuscript. C.-L.H., S.P.Y., W.J. and N.K.W. revised the manuscript. All authors approved the manuscript.

