## Supplementary material for "CNVPipe: An enhanced pipeline for accurate analysis of copy number variation from whole-genome sequencing": Suppl Figures and Tables

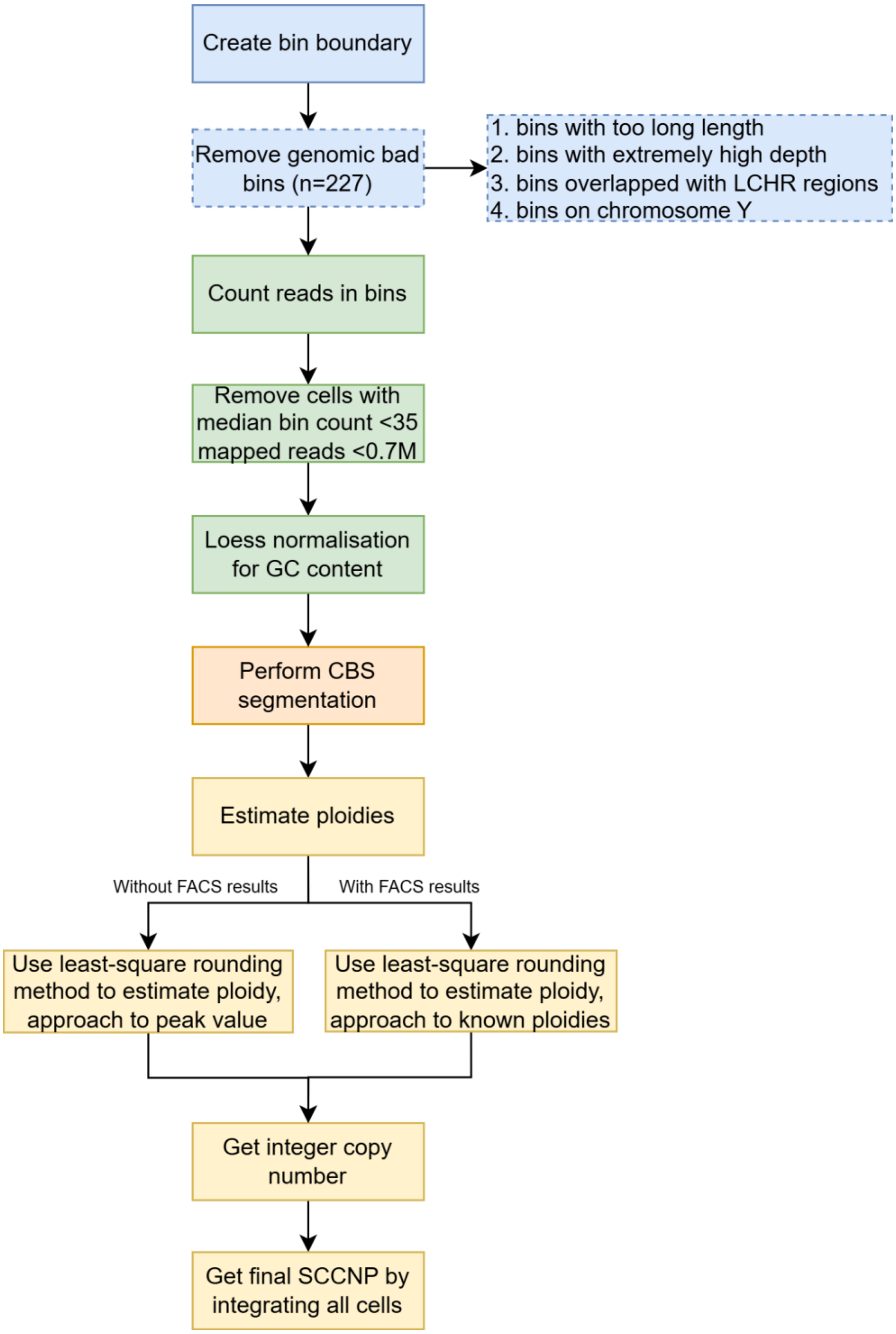


**Supplementary Figure 1. Detailed workflow of single-cell CNV calling.** CNVPipe single-cell module starts by creating a bin boundary file, dividing the whole human genome into 11,773 bins that have same number of mappable positions. Followed by counting the reads in each bin for single cells. Cells with low read depth will be filtered. After GC content correction by Loess normalization, genomic bad bins with abnormal length, read depth or overlapped with low-complexity high-repetitive regions will be removed. Circular binary segmentation method will be used to segment bins. Depending on whether the FACS data of single cell is accessible, the ploidy of single cancer cell is approached to known FACS peaks or inferred density peak after least-square rounding approach. Finally, all single-cell copy-number profiles will be combined together to generate a single-cell copy-number matrix.

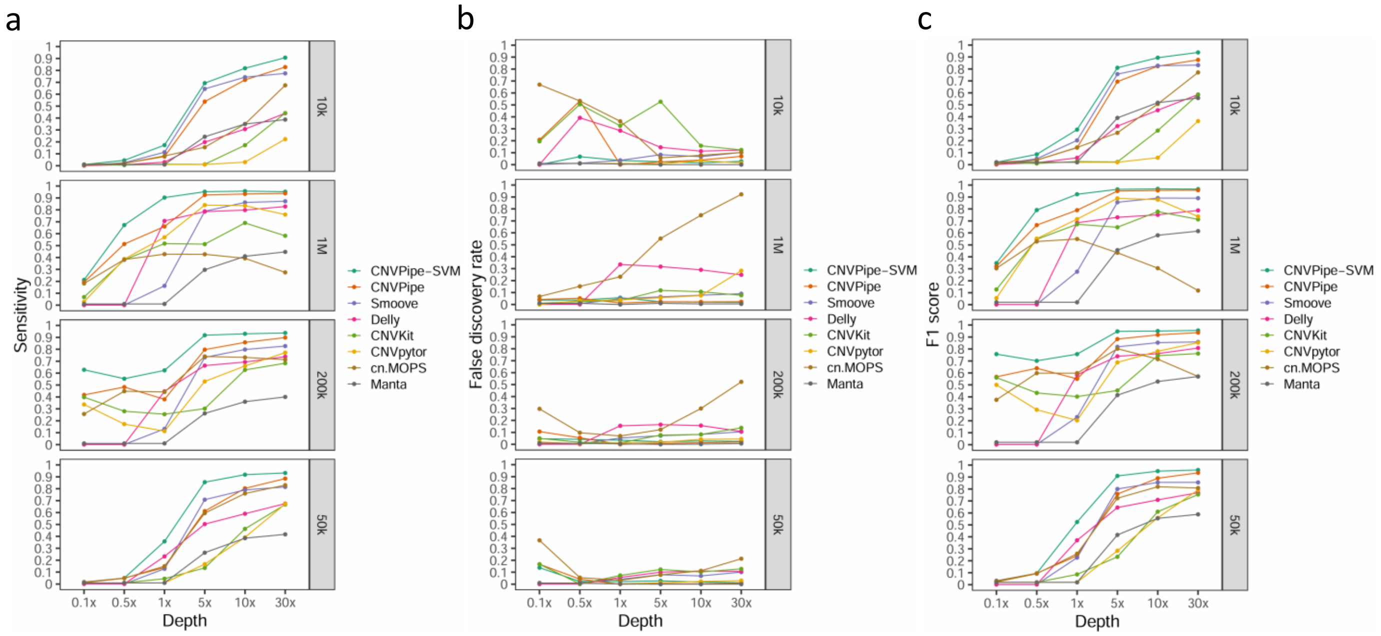


**Supplementary Figure 2. Performance evaluation of CNVPipe and individual CNV calling tools on simulation data with different CNV size and varying sequencing depth.** Line plot showing the averaged **a** sensitivity; **b** false discovery rate and **c** F1 score of different CNV calling tools on synthetic data. The X-axis represents different sequencing depth, from left to right, 0.1×, 0.5×, 1×, 5×, 10× and 30×, respectively. The facets represent groups with different CNV size, from top to bottom, 10 kb, 50 kb, 200 kb and 1 Mb. The colours represent different CNV calling tools including CNVPipe, Smoove, Delly, CNVKit, CNVpytor, cn.MOPS and Manta.

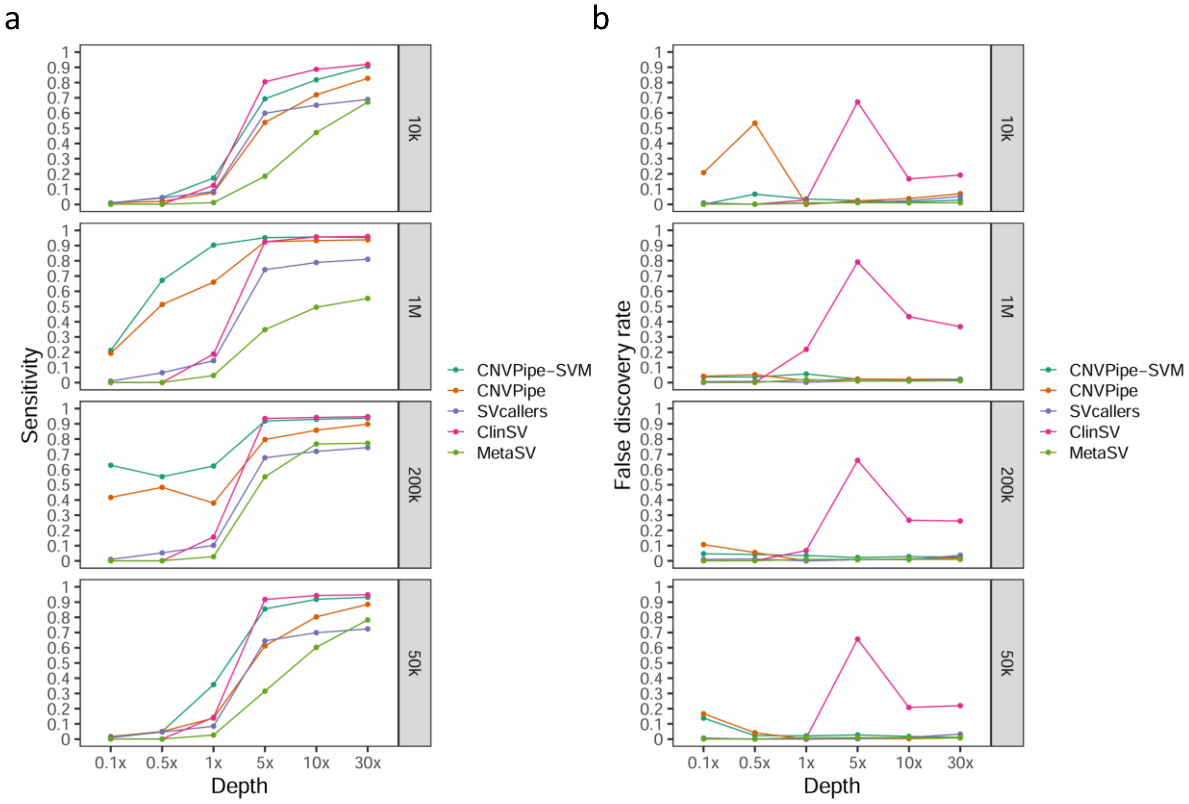


**Supplementary Figure 3. Performance evaluation of CNVPipe-SVM and ensemble CNV calling tools on simulation data with different CNV size and varying sequencing depth.** Line plot showing the averaged **a** sensitivity; **b** false discovery rate of different CNV calling tools on synthetic data. The X-axis represents different sequencing depth, from left to right, 0.1×, 0.5×, 1×, 5×, 10× and 30×, respectively. The facets represent groups with different CNV size, from top to bottom, 10 kb, 50 kb, 200 kb and 1 Mb. The colours represent different CNV calling tools including CNVPipe-SVM, CNVPipe, SVcallers, ClinSV and MetaSV.

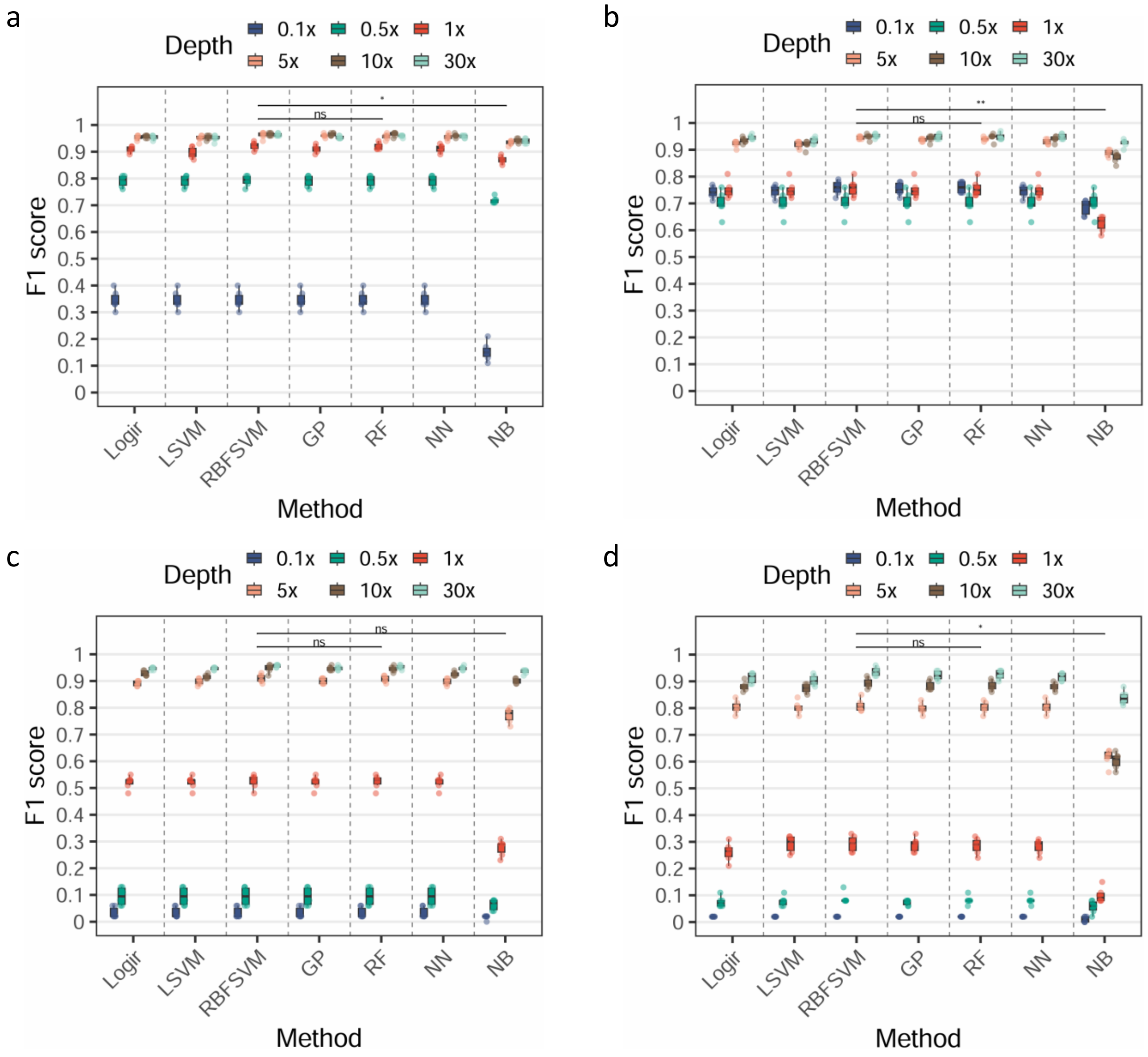


**Supplementary Figure 4. Boxplot of F1 score with statistical comparisons among 7 machine learning classifiers for simulated datasets with different CNV sizes.** The mean size of CNV is simulated to be **a** 1 million bases (1 Mb); **b** 200,000 base pairs (200 kb); **c** 50,000 base pairs (50 kb); and **d** 10,000 base pairs (10 kb). For each machine learning classifier, from leftmost box to rightmost box are sequencing depth of 0.1×, 0.5×, 1×, 5×, 10× and 30×. Logir: Logistic Regression; LSVM: Linear Support Vector Machine; RBFSVM: Radial Basis Function Support Vector Machine; GP: Gaussian Process; RF: Random Forest; NN: Neural Network; NB: Naïve Bayes. Wilcoxon test was performed between RBFSVM, RF and NB, * $p < 0.05$, ** $p < 0.01$, *** $p < 0.001$, ns: not significant.

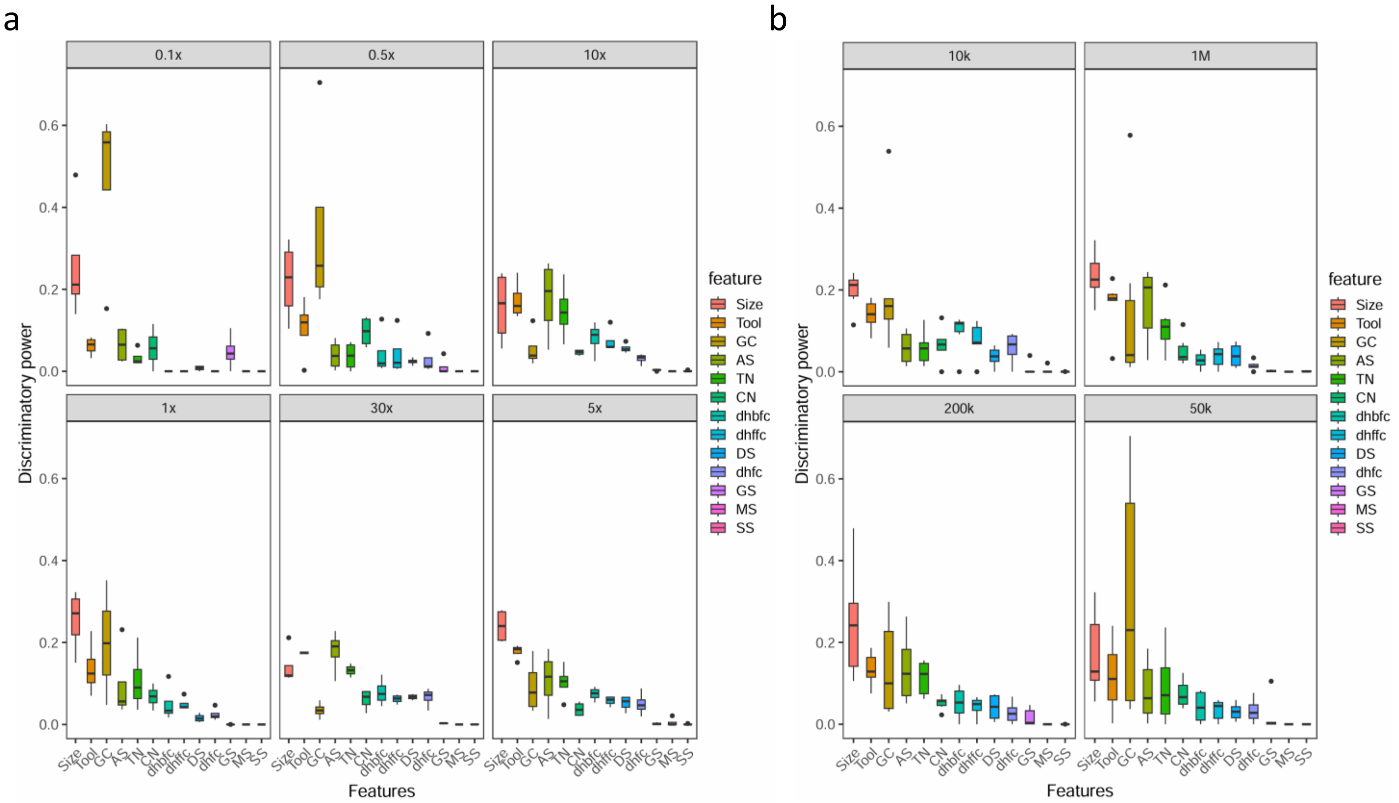


**Supplementary Figure 5. Boxplot of discriminatory power of 13 features in CNVPipe-SVM on simulated datasets. a** Each facet represents a sequencing depth; **b** each facet represents a CNV size. For each facet, the boxes from leftmost to rightmost are tool (CNV callers), tool number, CNV size, accumulated score, GC content, good score, depth score, snp score, copy number, mappability score, dhfc, dhbfc and dhffc.

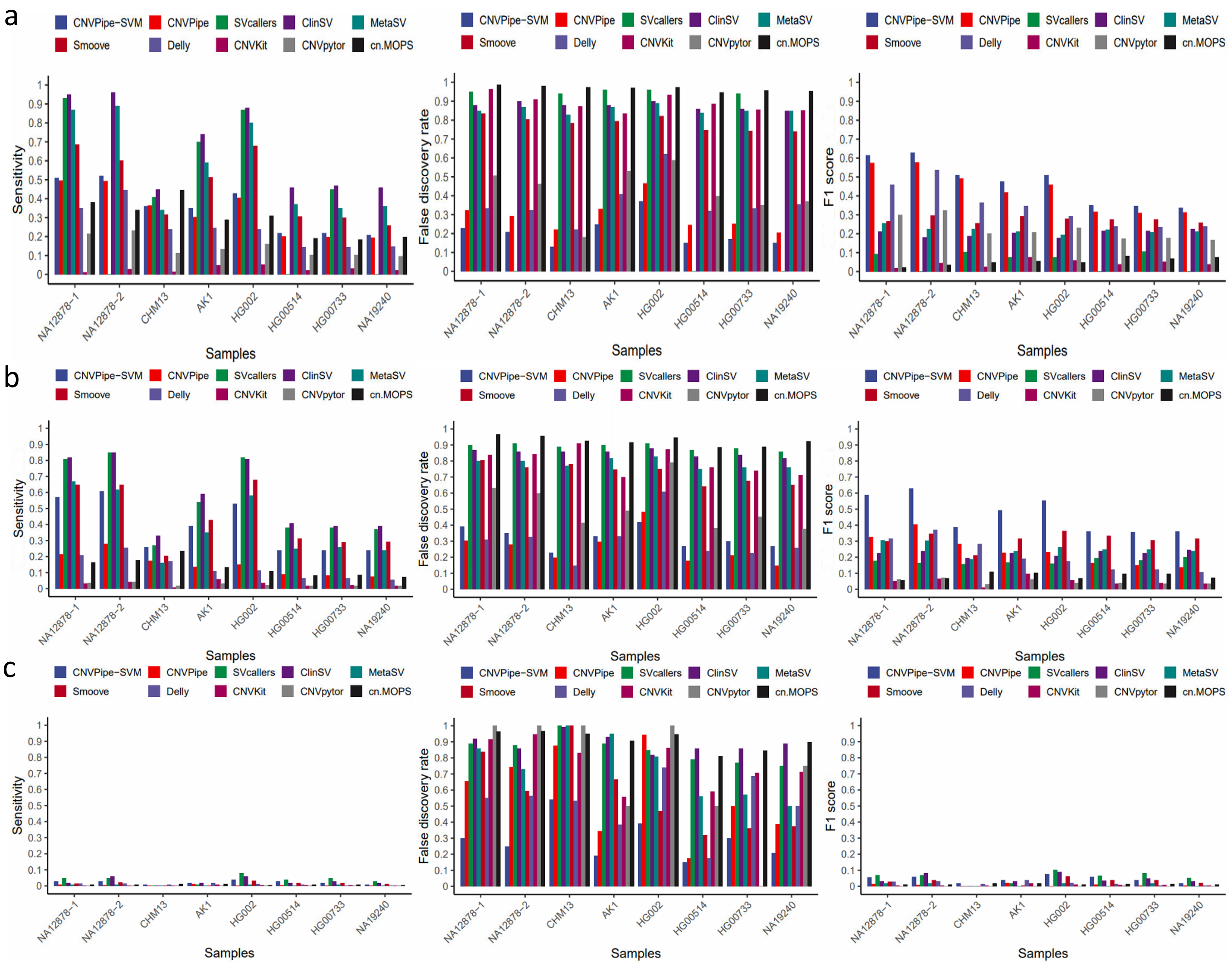


**Supplementary Figure 6. Sensitivity, false discovery rate and F1 score of ensemble and individual CNV callers on 8 authentic datasets.** Bar plot showing the sensitivity (left column), false discovery rate (middle column) and F1 score (right column) of CNV callers at sequencing depth of **a** 30×; **b** 10×; and **c** 1×, with X-axis represents authentic WGS samples, and Y-axis represents values of metrices. The Y-axis scale in all the panels are the same.

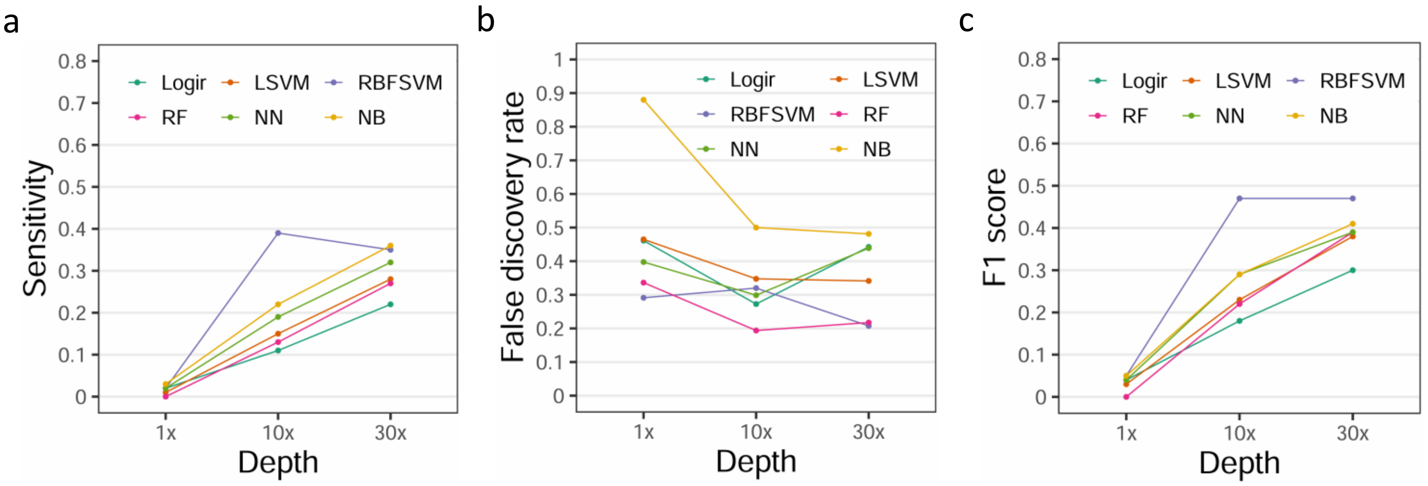


**Supplementary Figure 7. Line plot showing the performance of 6 machine learning classifiers on authentic datasets.** Average **a** sensitivity; **b** false discovery rate; and **c** F1 score of 6 classifiers on 8 authentic WGS samples under sequencing depth of 1×, 10× and 30×. Logir: Logistic Regression; LSVM: Linear Support Vector Machine; RBFSVM: Radial Basis Function Support Vector Machine; GP: Gaussian Process; RF: Random Forest; NN: Neural Network; NB: Naïve Bayes.

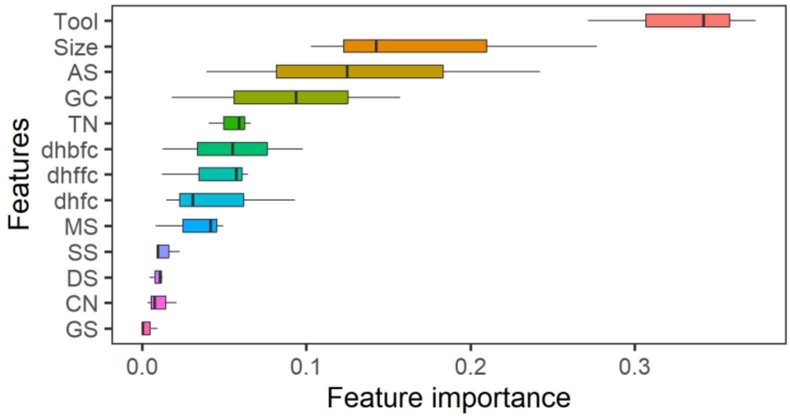


**Supplementary Figure 8. Boxplot of discriminatory power of 13 features in CNVPipe-SVM on real WGS datasets.** The boxes from top to bottom are tool (CNV callers), CNV size, accumulated score, GC content, tool number, dhbfc, dhffc and dhfc, mappability score, snp score, depth score, copy number and good score.

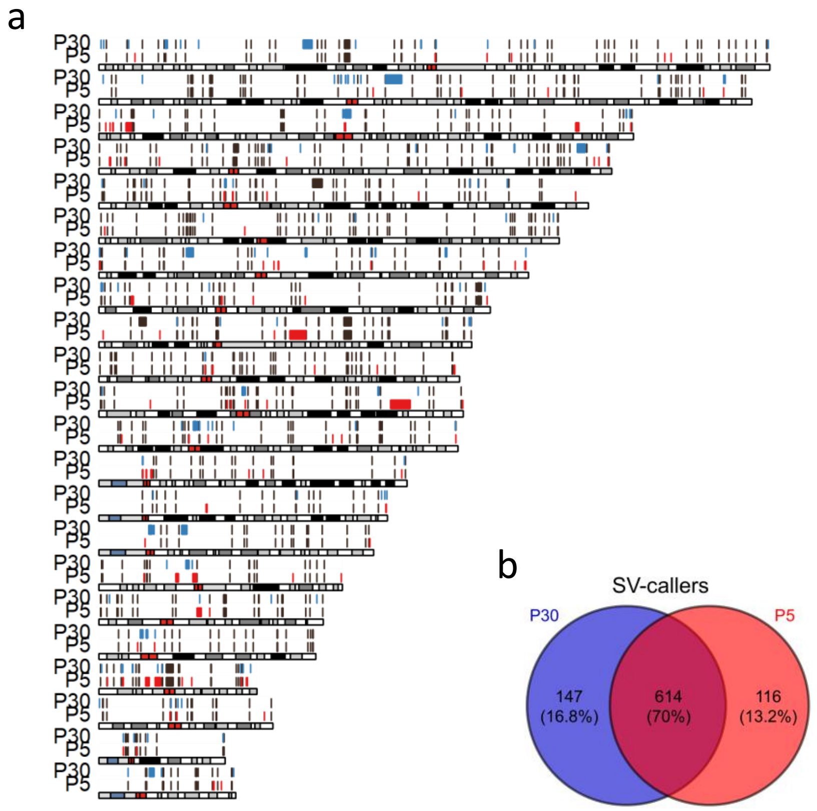


**Supplementary Figure 9. Distribution of CNVs identified in P30 and P5 by sv-callers. a** Karyoplot showing genomic position of CNVs identified in P30 and P5, green represents P30 unique CNVs, red represents P5 unique CNVs, and brown represents common CNVs between P30 and P5. **b** Percentage of unique and common CNVs in P30 and P5.

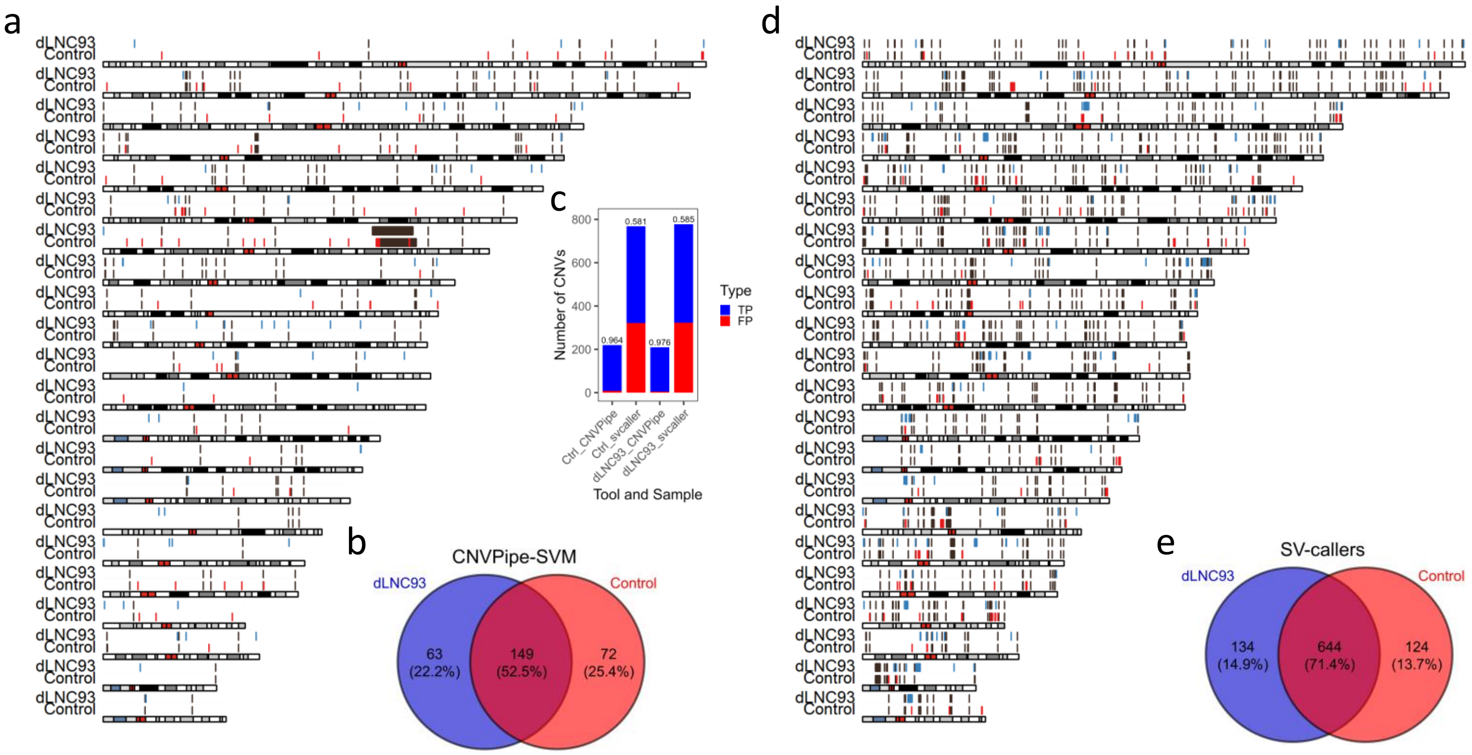


**Supplementary Figure 10. Distribution of CNVs identified in dLNC93 and CRISPR control by CNVPipe-SVM and sv-callers. a** Karyoplot showing genomic position of CNVs identified in dLNC93 and CRISPR control by CNVPipe-SVM. **b** Percentage of unique and common CNVs in dLNC93 and CRISPR control. **c** Percentage of true positive CNVs from CNVPipe-SVM and sv-callers manually inspected by CNView. **d** CNVs identified by sv-callers. **e** Percentage of unique and common CNVs in dLNC93 and CRISPR control identified by sv-callers. Green represents dLNC93 unique CNVs, red represents CRISPR control unique CNVs, and brown represents common CNVs between dLNC93 and CRISPR control.

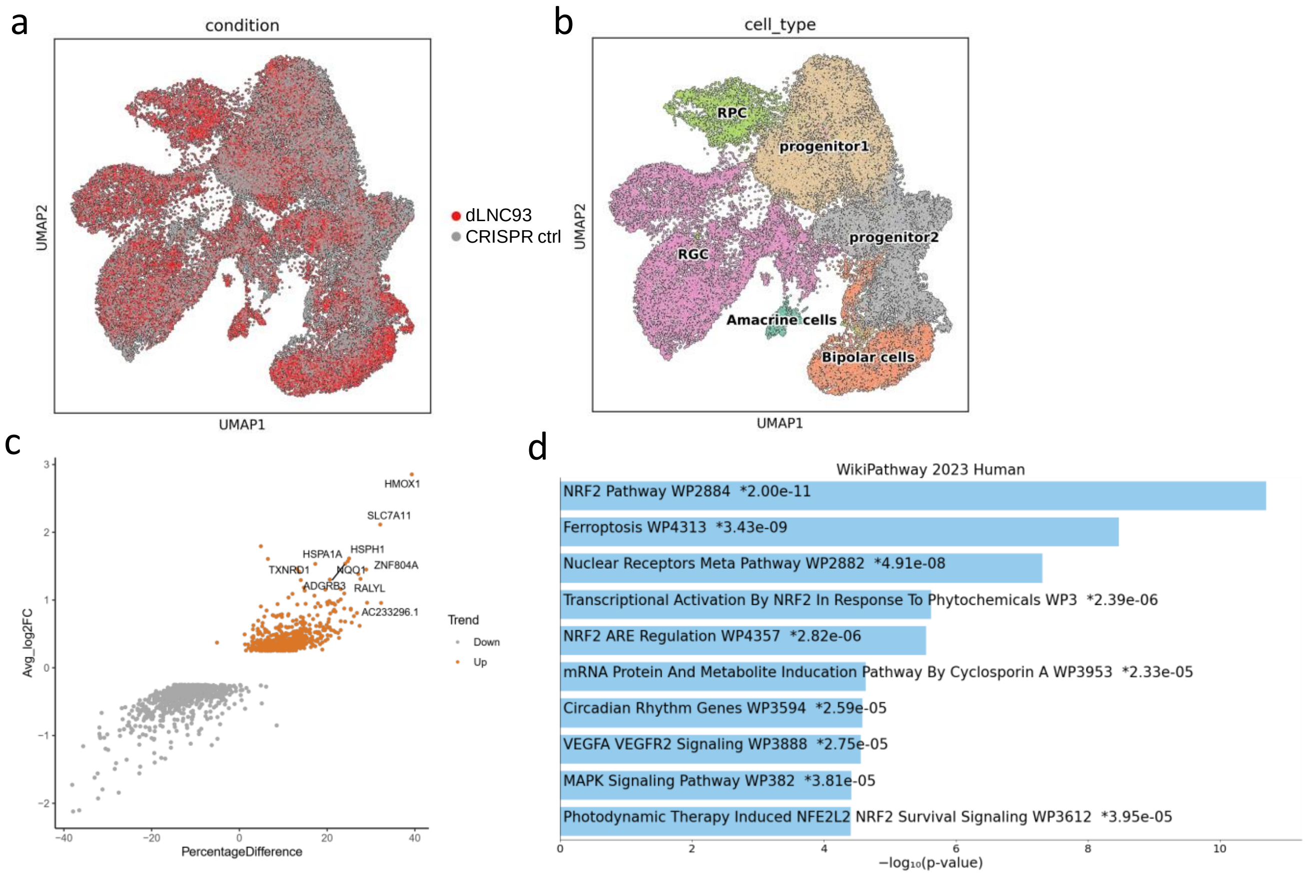


**Supplementary Figure 11. Single-cell RNA sequencing analysis of dLNC93 and control samples. a** UMAP plot of single cells differentiated from iPSC dLNC93 and control samples. **b** UMAP plot showing the annotation of cell types, RGC is identified and compared between dLNC93 and control sample. **c** Dotplot showing up-regulated genes in dLNC93 sample compared with control sample. **d** Enriched pathway of up-regulated genes. RGC: retinal ganglion cell; RPC: retinal progenitor cell.

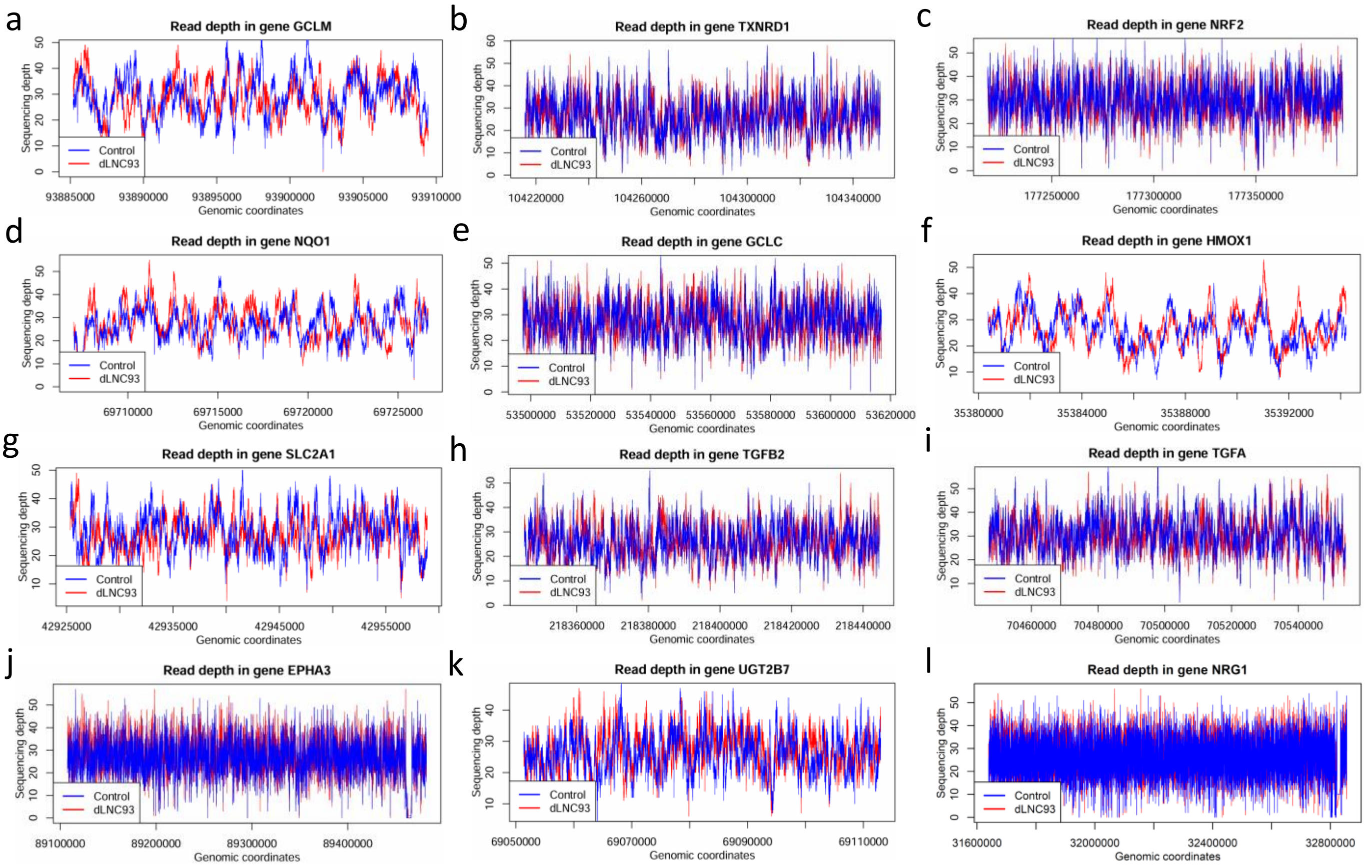


**Supplementary Figure 12. Comparison of read depth in six upregulated DEGs between dLNC93 and control sample. a** GCLM; **b** TXNRD1; **c** NRF2; **d** NQO1; **e** GCLC; **f** HMOX1; **g** SLC2A1; **h** TGFB2; **i** TGFA; **j** EPHA3; **k** UGT2B7; **l** NRG1.

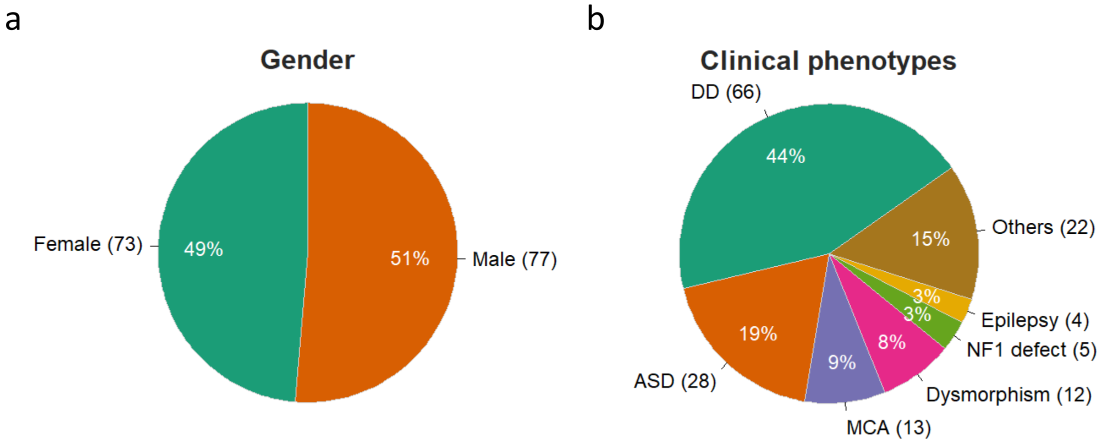


**Supplementary Figure 13. a** gender and **b** clinical phenotypes of 150 pediatric patients. DD: developmental delay; ASD: autism spectrum disorder; MCA: multiple congenital anomalies; others: other developmental defects.

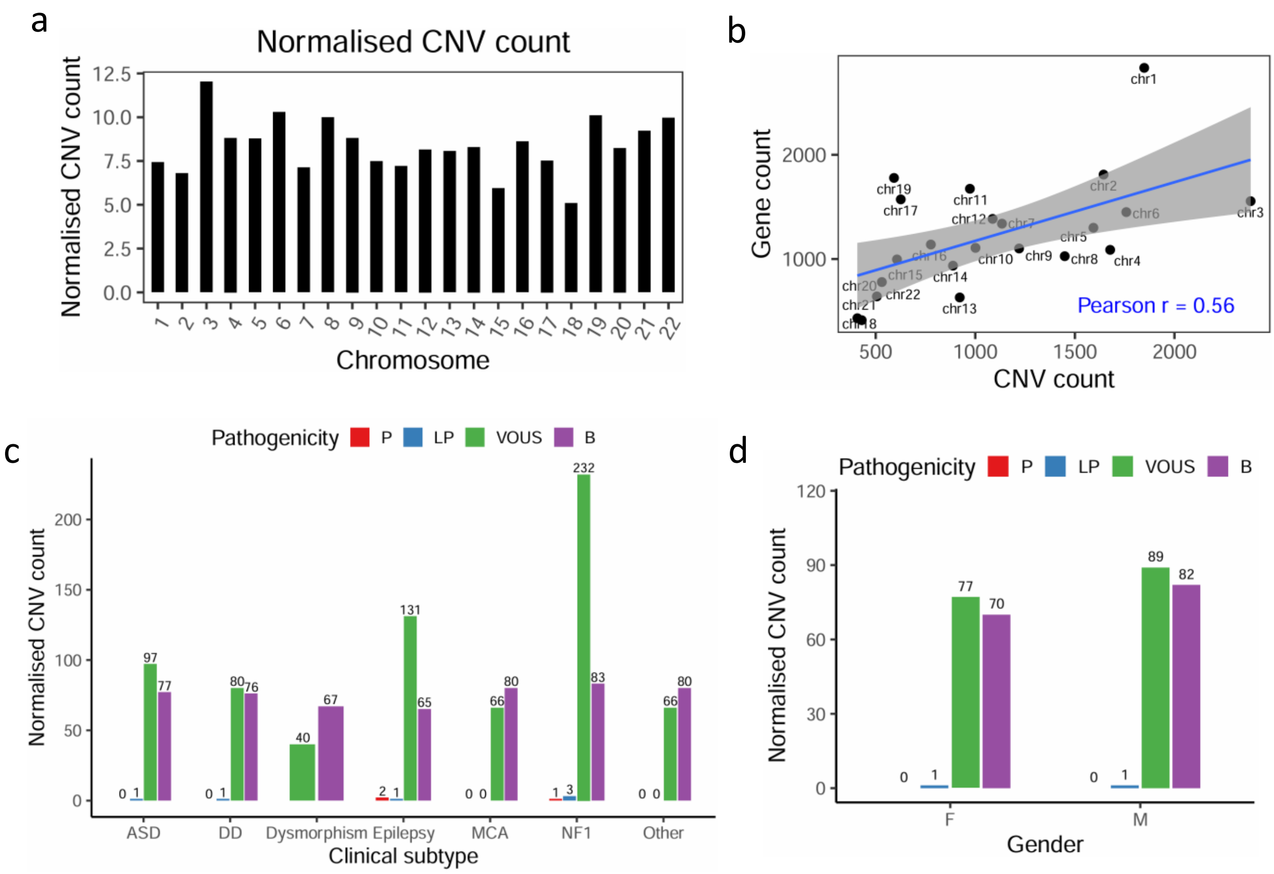


**Supplementary Figure 14. a** CNV count in chromosomes normalised by their length; **b** Correlation between CNV count and number of genes in chromosomes; **c** CNVs with known pathogenicity in different clinical subtypes normalized with patient number; **d** CNVs with known pathogenicity in different genders normalized with patient number.

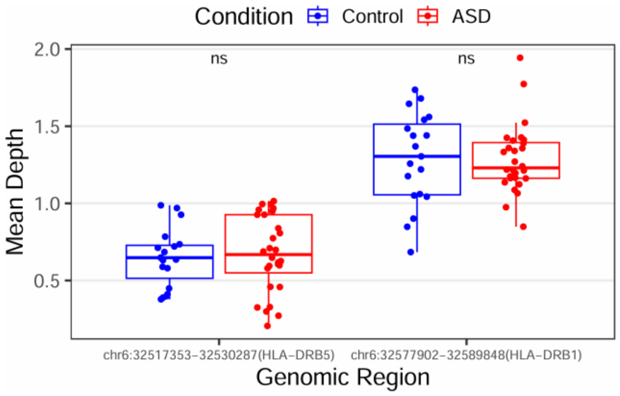


**Supplementary Figure 15. Mean coverage depth of ASD and control samples in HLA-DRB1 and HLA-DRB5 regions.** ns: not significant.

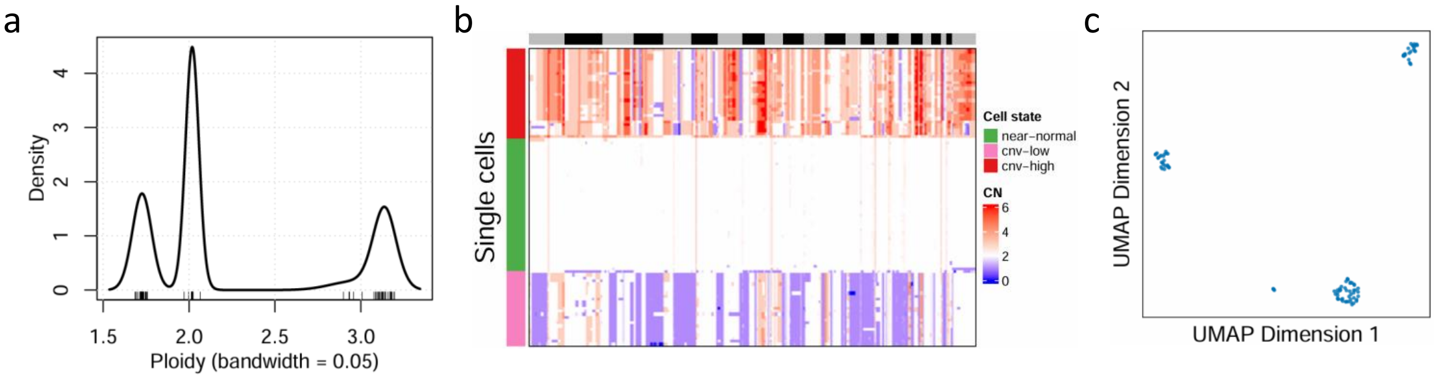


**Supplementary Figure 16. Results from using known ploidies for calculation. a** Density plot showing the ploidies calculated from final integer copy-number matrix; **b** Heatmap showing the whole-genome distribution of CNVs in all single cells; **c** UMAP plot showing the clustering of single cells based on integer copy number profile.

**Supplementary Table 1.** CNVs uniquely identified in P30 and P5 by CNVPipe-SVM

| Chromosome | Start | End | Length | CN in P30 | CN in P5 |
|---|---|---|---|---|---|
| | | **P30 Unique CNVs** | | | |
| chr1 | 120699900 | 120700100 | 200 | 4 | 3 |
| chr2 | 104819001 | 104819389 | 388 | 3 | 2 |
| chr2 | 121617298 | 121618399 | 1101 | 3 | 2 |
| chr4 | 48165100 | 48165800 | 700 | 4 | 2 |
| chr7 | 56891654 | 56892654 | 1000 | 1 | 2 |
| chr10 | 83762959 | 83764167 | 1208 | 0 | 1 |
| chr12 | 8411302 | 8411602 | 300 | 4 | 2 |
| chr13 | 113234999 | 113235199 | 200 | 3 | 2 |
| chr16 | 89878000 | 89879200 | 1200 | 3 | 2 |
| chr19 | 475799 | 476199 | 400 | 4 | 2 |
| | | **P5 Unique CNVs** | | | |
| chr1 | 234780195 | 234781195 | 1000 | 2 | 4 |
| chr7 | 114829500 | 114829800 | 300 | 2 | 1 |
| chr8 | 43657600 | 43657750 | 150 | 2 | 3 |
| chr9 | 105057599 | 105057994 | 395 | 2 | 4 |

CN: copy number.

**Supplementary Table 2**. Unique CNVs in P30 and P5 validated by ddPCR

| Concentration (copies/ul) | | | | |
|---|---|---|---|---|
| CNV1 (P30U_chr1: 1206999000-120700100) | | | | |
| Sample | CNV1 | RPP30 | CN | CNVPipe output |
| P5 iPSC | 320 | 76 | 2.11 | 3 |
| P30 iPSC | 506 | 73.2 | 3.46 | 4 |
| CNV2 (P30U_chr2: 104819001-104819389) | | | | |
| Sample | CNV2 | RPP30 | CN | CNVPipe output |
| P5 iPSC | 72 | 76 | 1.89 | 2 |
| P30 iPSC | 137 | 73.2 | 3.74 | 3 |
| CNV3 (P30U_chr2: 121617298-121618399) | | | | |
| Sample | CNV3 | RPP30 | CN | CNVPipe output |
| P5 iPSC | 72 | 76 | 1.89 | 2 |
| P30 iPSC | 137 | 73.2 | 3.74 | 3 |
| CNV4 (P30U_chr4: 48165100-48165800) | | | | |
| Sample | CNV4 | RPP30 | CN | CNVPipe output |
| P5 iPSC | 79.3 | 76 | 2.09 | 2 |
| P30 iPSC | 143 | 73.2 | 3.91 | 4 |
| CNV5 (P30U_chr7: 56891654-56892654) | | | | |
| Sample | CNV5 | RPP30 | CN | CNVPipe output |
| P5 iPSC | 150 | 76 | 1.97 | 2 |
| P30 iPSC | 92.5 | 73.2 | 1.26 | 1 |
| CNV6 (P30U_chr10: 83762959-83764167) | | | | |
| Sample | CNV6 | RPP30 | CN | CNVPipe output |
| P5 iPSC | 31.2 | 76 | 0.82 | 1 |
| P30 iPSC | 7 | 73.2 | 0.19 | 0 |
| CNV7 (P30U_chr12: 8411302-8411602) | | | | |
| Sample | CNV7 | RPP30 | CN | CNVPipe output |
| P5 iPSC | 69.2 | 76 | 1.82 | 2 |
| P30 iPSC | 139.7 | 73.2 | 3.82 | 4 |
| CNV8 (P30U_chr13: 113234999-113235199) | | | | |
| Sample | CNV8 | RPP30 | CN | CNVPipe output |
| P5 iPSC | 84.5 | 76 | 2.22 | 2 |
| P30 iPSC | 141 | 73.2 | 3.85 | 3 |
| CNV9 (P30U_chr16: 89878000-89879200) | | | | |
| Sample | CNV9 | RPP30 | CN | CNVPipe output |

| | | | | |
|---|---|---|---|---|
| P5 iPSC | 75.2 | 76 | 1.98 | 2 |
| P30 iPSC | 127 | 73.2 | 3.47 | 3 |
| CNV10 (P30U_chr19: 475799-476199) | | | | |
| Sample | CNV10 | RPP30 | CN | CNVPipe output |
| P5 iPSC | 78 | 76 | 2.05 | 2 |
| P30 iPSC | 169 | 73.2 | 4.62 | 4 |
| CNV11 (P5U_chr1: 247118900-247119300) | | | | |
| Sample | CNV11 | RPP30 | CN | CNVPipe output |
| P5 iPSC | 157 | 76 | 4.13 | 4 |
| P30 iPSC | 59.3 | 73.2 | 1.62 | 2 |
| CNV12 (P5U_chr7: 114829500-114829800) | | | | |
| Sample | CNV12 | RPP30 | CN | CNVPipe output |
| P5 iPSC | 51.8 | 76 | 1.36 | 1 |
| P30 iPSC | 73 | 73.2 | 1.99 | 2 |
| CNV13 (P5U_chr9_105057599-105057994) | | | | |
| Sample | CNV13 | RPP30 | CN | CNVPipe output |
| P5 iPSC | 174 | 76 | 4.58 | 4 |
| P30 iPSC | 78.6 | 73.2 | 2.15 | 2 |
| CNV14 (93U_chr6: 76726700-76727000) | | | | |
| Sample | CNV14 | RPP30 | CN | CNVPipe output |
| CRISPR ctrl iPSC | 44.3 | 82 | 1.08 | 1 |
| dLNC93 iPSC | 115 | 78.4 | 2.93 | 4 |
| CNV15 (93U_chr11: 107919500-107919700) | | | | |
| Sample | CNV15 | RPP30 | CN | CNVPipe output |
| CRISPR ctrl iPSC | 83.5 | 82 | 2.04 | 2 |
| dLNC93 iPSC | 36.1 | 78.4 | 0.92 | 1 |
| CNV16 (CtrlU_chr1: 247118900-247119300) | | | | |
| Sample | CNV16 | RPP30 | CN | CNVPipe output |
| CRISPR ctrl iPSC | 165 | 82 | 4.02 | 4 |
| dLNC93 iPSC | 95.5 | 78.4 | 2.44 | 2 |
| CNV17 (CtrlU_chr13: 18910000-18910200) | | | | |
| Sample | CNV17 | RPP30 | CN | CNVPipe output |
| CRISPR ctrl iPSC | 128 | 82 | 3.12 | 3 |
| dLNC93 iPSC | 73.6 | 78.4 | 1.88 | 2 |

**Supplementary Table 3**. CNVs identified in ASD patients overlapped with ASD-risk genes

| Chromosome | Start | End | Samples | Genes |
|---|---|---|---|---|
| chr1 | 10630034 | 10658249 | 27859, 28102, 28448, 28941, 29683, 29849 | CASZ1 |
| chr1 | 151360573 | 151438099 | 12965, 20469, 29355, 29463, 30411 | POGZ |
| chr2 | 25330101 | 25346507 | 21866, 27859, 29540, 29888 | DNMT3A |
| chr3 | 48409601 | 48418915 | 20195, 28102, 29405, 29540, 29683, 29960, 30610, 30815 | PLXNB1 |
| chr3 | 71433230 | 71442599 | 20195, 29405, 29540, 29615, 29746, 29888, 29960 | FOXP1 |
| chr3 | 177189801 | 177205299 | 20195, 21866, 28941, 29195, 29540, 29615, 29746, 29849, 29888, 29960, 30610, 30815 | TBL1XR1 |
| chr5 | 14139101 | 14150027 | 20195, 21866,27859, 28448, 28941, 29405, 29540, 29595, 29615, 29683, 29746, 29849, 30610, 30815 | TRIO |
| chr5 | 14477001 | 14498699 | 20195, 27859, 28448, 29595, 30610 | TRIO |
| chr6 | 156770101 | 156788699 | 20195, 21866, 27859, 28102, 28448, 28941, 29195, 29405, 29540, 29595, 29683, 29746, 29849, 29888, 29960, 30815 | ARID1B |
| chr6 | 157145201 | 157151399 | 20195, 29405, 29540, 29595, 29615, 29849, 29888, 29960, 30610, 30815 | ARID1B |
| chr9 | 137715283 | 137726799 | 19097, 29195, 29746, 30815 | EHMT1 |
| chr10 | 112945401 | 112960899 | 27859, 28102, 28448, 29405 | TCF7L2 |
| chr10 | 129957576 | 129977540 | 27859, 28448, 29849, ~~30237~~ | EBF3 |
| chr12 | 13847701 | 13860099 | 28102, 29405, 29746, 29888, 29960 | GRIN2B |
| chr12 | 56181301 | 56196799 | 21866, 29195, 29615, 29746, 29888, 30815 | SMARCC2 |
| chr17 | 31046501 | 31948599 | 19097, 24975, 25169, 30361 | NF1 |
| chr18 | 33568557 | 33591599 | 28102, 29595, 29888, 29960 | ASXL3 |
| chr22 | 50669501 | 50725299 | 20195, 27859, 28448, 29195, 30610 | SHANK3 |

**Supplementary Table 4**. Recurrent CNVs appeared in over three ASD patients

| Chr | Start | End | CN | Samples | Pathogenicity |
|---|---|---|---|---|---|
| chr2 | 79100750 | 79118199 | 4 | 26503,26475,27859 | Uncertain |
| chr4 | 62802901 | 62812199 | 0 | 30835,26476,29460,30587 | Uncertain |
| chr9 | 101952331 | 101965199 | 0 | 30213,22445,28941,29405 | Benign |
| chr10 | 46512289 | 46561999 | 4 | 22445,26792,30587 | Uncertain |
| chr15 | 99875801 | 99885271 | 0 | 29265,26672,30171,26565 | Benign |
| chr17 | 15139959 | 15157728 | 0 | 22445,29601,26565,27859,30171,30078 | Uncertain |
| chr19 | 27338901 | 27400899 | 6 | 30078,26475,26505 | Uncertain |
| chr21 | 7915746 | 8049838 | 4 | 29264,26505,29601 | Uncertain |

**Supplementary Table 5**. Transcription factor binding sites overlapped with recurrent CNVs in ASD patients

| Overlapped TFBS of | Chromosome | Start | End | Length (bp) |
|---|---|---|---|---|
| POLR2A | chr2 | 79115024 | 79115742 | 718 |
| ZNF263 | chr2 | 79114919 | 79115503 | 584 |
| CTCF | chr2 | 79105056 | 79105896 | 840 |
| FOXA1 | chr4 | 62806902 | 62807138 | 236 |
| ESR1 | chr4 | 62807023 | 62807293 | 270 |
| GATA3 | chr9 | 101957711 | 101958187 | 476 |
| REST | chr9 | 101955040 | 101955450 | 410 |
| MAX | chr17 | 15143548 | 15143944 | 396 |
| CEBPB | chr17 | 15154096 | 15154352 | 256 |
| RFX1 | chr19 | 27365590 | 27365990 | 400 |

**Supplementary Table 6**. Candidate cis-regulatory elements overlapped with recurrent CNVs in ASD patients

| Chromosome | Start | End | EncodeLabel | Length |
|---|---|---|---|---|
| chr2 | 79103304 | 79103554 | dELS | 250 |
| chr4 | 62806582 | 62806744 | dELS | 162 |
| chr4 | 62807416 | 62807755 | dELS | 339 |
| chr4 | 62807759 | 62807972 | dELS | 213 |
| chr9 | 101957345 | 101957505 | dELS | 160 |
| chr9 | 101957767 | 101958104 | dELS | 337 |
| chr10 | 46516676 | 46517026 | dELS | 350 |
| chr10 | 46547119 | 46547320 | dELS | 201 |
| chr10 | 46547763 | 46548112 | dELS | 349 |
| chr10 | 46548196 | 46548348 | dELS | 152 |
| chr10 | 46554519 | 46554720 | pELS | 201 |
| chr10 | 46555403 | 46555620 | PLS | 217 |
| chr10 | 46555844 | 46556002 | pELS | 158 |
| chr15 | 99878894 | 99879200 | dELS | 306 |

**Supplementary Table 7**. ddPCR results validating TFBS overlapped with recurrent CNVs

| Concentration (copies/ul) | | | | |
|---|---|---|---|---|
| Sample | POLR2A chr2_79115024_79115742 | RPP30 | CN | CNVPipe output |
| 26475 | 342 | 85.3 | 8.02 | 3 |
| 26503 | 652 | 132 | 9.88 | 4 |
| 27859 | 16 | 15.7 | 2.04 | 3 |
| C1 | 291 | 116 | 5.02 | |
| | CTCF chr2_79105056_79105896 | | | |
| 26475 | 376 | 85.3 | 8.82 | 3 |
| 26503 | 1441 | 132 | 21.83 | 4 |
| 27859 | 25.7 | 15.7 | 3.27 | 3 |
| C1 | 240 | 116 | 4.14 | |
| | FOXA1 chr4_62806902_62807138 | | | |
| 26476 | 0.13 | 69 | 0.00 | 0 |
| 29460 | 0 | 87.2 | 0.00 | 0 |
| 30587 | 0 | 94 | 0.00 | 0 |
| 30835 | 52.2 | 104.1 | 1.00 | 0 |
| C1 | 0.21 | 116 | 0.00 | |
| | ESR1 chr4_62807023_62807293 | | | |
| 26476 | 1.7 | 69 | 0.05 | 0 |
| 29460 | 1.45 | 87.2 | 0.03 | 0 |
| 30587 | 1.6 | 94 | 0.03 | 0 |
| 30835 | 52.7 | 104.1 | 1.01 | 0 |
| C1 | 1.21 | 116 | 0.02 | |
| | GATA3 chr9_101957711_101958187 | | | |
| 22445 | 0 | 124 | 0.00 | 0 |
| 28941 | 119 | 111 | 2.14 | 0 |
| 29405 | 0.22 | 112 | 0.00 | 0 |
| 30213 | 0.06 | 121 | 0.00 | 0 |
| C1 | 117 | 116 | 2.02 | |
| Sample | REST chr9_101955040_101955450 | | | |

| | | | | |
|---|---|---|---|---|
| 22445 | 0 | 124 | 0.00 | 0 |
| 28941 | 144 | 111 | 2.59 | 0 |
| 29405 | 0 | 112 | 0.00 | 0 |
| 30213 | 0 | 121 | 0.00 | 0 |
| C1 | 130 | 116 | 2.24 | |
| | MAX chr17_15143548_15143944 | | | |
| 22445 | 7 | 191 | 0.07 | 0 |
| 26565 | 4 | 267 | 0.03 | 0 |
| 27859 | 4.8 | 15.7 | 0.61 | 0 |
| 29601 | 7.2 | 216 | 0.07 | 1 |
| 30078 | 144 | 195 | 1.48 | 1 |
| 30171 | 123 | 191 | 1.29 | 1 |
| C1 | 453 | 491 | 1.85 | |
| | CEBPB chr17_15154096_15154352 | | | |
| 22445 | 0.13 | 191 | 0.00 | 0 |
| 26565 | 0.07 | 267 | 0.00 | 0 |
| 27859 | 9.2 | 15.7 | 1.17 | 0 |
| 29601 | 0 | 216 | 0.00 | 1 |
| 30078 | 236 | 195 | 2.42 | 1 |
| 30171 | 157 | 191 | 1.64 | 1 |
| C1 | 435 | 491 | 1.77 | |
| | RFX1 chr19_27365590_27365990 | | | |
| 26475 | 176 | 77.3 | 4.55 | 3 |
| 26505 | 167 | 110 | 3.04 | 3 |
| 30078 | 154 | 75.4 | 4.08 | 3 |
| C1 | 249 | 116 | 4.29 | |

**Supplementary Table 8**. Recommended calling resolutions for different depth data

| Depth of data | Resolutions |
|---|---|
| 0.1x | 369300 |
| 0.5x | 73800 |
| 1x | 36800 |
| 5x | 7400 |
| 10x | 3600 |
| 30x | 1200 |

**Supplementary Table 9**. Information of real WGS benchmark set

| ID | NGS info | Reads num | SV set resource | Mean size of CNV | Type | Ref. |
|---|---|---|---|---|---|---|
| NA12878-1 | 2x100bp | 1,436,823,773 | svclassify [1] | 4880.94 | DEL | [2] |
| NA12878-2 | 2x100bp | 787,265,109 | svclassify [1] | 4880.94 | DEL | [3] |
| CHM13 | 2x100bp | 456,931,559 | nstd137 | 3467.99 | DEL | [4] |
| AK1 | 2x150bp | 432,004,180 | original paper | 12734.5 | DEL | [5] |
| HG002 | 2x147bp | 268,996,648 | nstd175 | 4036.86 | DEL | [6] |
| HG00514 | 2x125bp | 519,884,509 | nstd152 | 15169.7 | DEL, DUP | [7] |
| HG00733 | 2x125bp | 411,722,389 | nstd152 | 13385.8 | DEL, DUP | [7] |
| NA19240 | 2x125bp | 521,786,371 | nstd152 | 13580.4 | DEL, DUP | [7] |
