## Supplementary material for "CNVPipe: An enhanced pipeline for accurate analysis of copy number variation from whole-genome sequencing": Suppl Methods

### 1. Parameters used in CNVPipe to run different CNV calling tools

#### 1.1 CNVKit

CNVKit divides the genome into fixed-size bins and counts read depth in those bins, followed by log2 transformation and centering. It applies a rolling median method to correct the biases caused by GC content and repetitive regions. Control samples could alternatively be provided to normalize the read depth and thus remove the CNVs that potentially exist in healthy samples. Before running CNVKit for CNV calling, we generated an accessibility file by 'access' command in CNVKit. The first excluded region file is obtained from UCSC, containing the genomic coordinates of centromeres, telomeres and heterochromatin. The second excluded region file is obtained from 10X genomics (http://cf.10xgenomics.com/supp/genome/GRCh38/sv_blacklist.bed), which contains high repetitive and low complexity regions in human genome. We called CNVs for all samples by CNVKit by 'batch' command, with '-m wgs' assigning whole-genome sequencing mode. Then we used 'segmetrics' command to calculate confidence interval of log2 ratio of segments. We then use 'call' command with '-m clonal –filter ci' to determine integer copy number from log2-transformed copy ratio of segments and merge segments with similar confidence interval. Finally, we wrote a Python script to convert .cns format into bed format, and to merge consecutive CNVs with identical type of CNVs but different copy numbers. The merged region will take rounded average copy number.

#### 1.2 CNVpytor

CNVpytor starts with counting read depth and correcting depth ratio in fixed-size bins, but differently, it uses a mean-shift approach to segment bins and uses statistical test like t-test to determine whether a segment is a reliable CNV region. CNVpytor does not require control samples and reference genome, neither exclude any bad regions in the genome. The usage of CNVpytor basically follows the developer's instructions on GitHub (https://github.com/abyzovlab/CNVpytor), that is: use '-rd' command to extract reads from bam file, use '-his' command to calculate reads depth at specified resolution, use '-partition' to segment bins by mean-shift method, and finally, use '-call' to call CNVs and calculate integer copy number. For each CNV, CNVpytor gives some statistics for us to do filtering: 'eval1' means e-value (p-value multiplied by genome size divided by bin size) calculated by t-test between RD statistics in the region and global; 'eval2' means e-value from the probability of RD values within the region to be in the tails of a gaussian distribution of binned RD; 'q0' means fraction of reads mapped with q0 quality in call region; 'pN' means fraction of reference genome gaps (Ns) in call region; 'dG' means distance from closest large (>100bp) gap in reference genome. We wrote a Python script to filter high-confidence CNVs by letting max (eval1, eval2) < 0.00001 and q0 < 0.5 and pN < 0.5 and dG > 10000.

#### 1.3 cn.MOPS

cn.MOPS calls CNVs based on assumptions that normalized read counts follow Poisson distribution among samples and the copy number of all bins is most likely to be 2. It uses a mixture of Poissons model to decompose the read depth in each bin across samples, and segments bins based on sI/NI call, detecting variations across samples and along chromosomes. It requires at least six samples to run as it uses a mixture of Poisson model to decompose the read depth among several samples. To run it, we firstly used the function 'getReadCountsFromBAM' in cn.MOPS R package to extract read counts at specified resolution, then we used the function 'cn.mops' followed by 'callIntegerCopyNumbers' to call CNVs and calculate integer copy number for all samples, finally, we transformed Granges objects into bed-like format and store them into file by samples. Just like for CNVKit results, we also wrote a Python Script to merge consecutive CNVs with identical type of CNVs but different copy numbers.

##### 1.4 Smoove

Smoove is a tool that wrapped Lumpy but greatly simplifies and speeds SV calling for short reads, it also integrates other tools like duphold and SVtyper to genotype or refine high-confidence CNVs. Lumpy starts with extracting discordant and split reads from mapped bam files, then apply local assembly to determine the breakpoints of variations, followed by a genotyping process to refine and extract high-confidence SVs. Traditionally, we need to extract discordant reads and split reads from bam files before using Lumpy to call SVs. But Smoove integrated all these pre-processing steps, so we only need to use the Smoove 'call' command to call variations in one step. We provided the 10X SV black list file as exclusion regions to tell Smoove avoiding calling SVs in those regions. Then we used bcftools to extract 'DEL' and 'DUP' CNVs from vcf files and convert them into bed file for easier downstream CNV merging. The 'DEL' CNVs were assigned copy number 1, while the 'DUP' CNVs were assigned copy number 3. Since Smoove might produce conflict results for some noisy regions or complex SVs, we wrote a Python script to resolve those conflicts and select CNVs with size larger than 1kb.

##### 1.5 Delly

Delly applies a two-step strategy to call SVs: in the first step, extracts discordant paired-end reads and determines a potential SV region by finding discordant reads cluster; In the second step, extracts all split reads around SVs identified in the first step, and performs consensus assembly and alignment to determine the final SVs with single nucleotide breakpoint resolution. To use Delly, we downloaded mappability files according to developer's instructions (https://gear.embl.de/data/delly/). The developer suggests to call SVs first by 'call' command, and then use the SV set as input to refine the CNV calling. However, in our primary trail, Delly SV calling is very time-consuming, it took around 24 hours to call SVs for a 10x-depth sample. Therefore, in CNVPipe, we skipped Delly SV calling step, and directly used 'cnv' command to call CNVs. Similar as in Smoove, we used bcftools to extract CNVs with 'PASS' signal from vcf files and convert them into bed file.

#### 2. Parameters used in CNVPipe to do SNP calling

##### 2.1 GATK4 HaplotypeCaller

We basically followed the GATK best practice of germline SNP calling for single sample, that is: firstly, perform base quality recalibration as we mentioned in pre-processing step, then use 'HaplotypeCaller' module to get raw variant file, followed by 'VariantRecalibrator' and 'ApplyVQSR' to recalibrate the variant quality and select only SNPs.

##### 2.2 Freebayes

We used Freebayes to call SNPs for median-depth data, only recalibrated bam file and reference genome file are required for Freebayes to run. After getting the raw variant file, we used bcftools to do hard filtering with the condition of "-s LOWQUAL -e 'QUAL<10 || FMT/DP <5' --SnpGap 5", then also used bcftools to specifically extract SNPs to get final variant file.

#### 3. Scoring metrics

##### 3.1 Accumulative score

During iterative CNV merging, if two CNVs are found to have overlaps, the length of overlapped region is firstly calculated, then it is divided by the length of the merged CNV. We multiply the accumulated overlap fractions by 100 and regard it as the accumulative score.

##### 3.2 Depth score

Duphold calculates DHFFC and DHBFC for each merged CNVs. DHFFC represents fold-change for the variant depth relative to flanking regions, while DHBFC presents fold-change for the variant depth relative to bins in the genome with similar GC-content. For copy number deletions, they will be assigned 100 points if DHFFC $< 0.5$, 80 points if $0.5 <$ DHFFC $< 0.7$, 50 points if $0.7 <$ DHFFC $< 0.99$ and 0 point otherwise. For copy number duplications, they will be assigned 100 points if DHBFC $> 1.5$, 80 points if $1.3 <$ DHBFC $< 1.5$, 50 points if $1.01 <$ DHBFC $< 1.3$, and 0 point otherwise.

##### 3.3 SNP support score

CNVfilteR calculates the number of total variants, heterozygous variants and homozygous variants in CNV regions, and assigns confidence scores for deletion and duplication by the number of homozygous variants or B-allele frequency of heterozygous variants. In CNVPipe, we only use the final output of CNVfilteR as well as ‘True’ or ‘False’ labelling, thus ‘True’ CNVs will be assigned 100 points, while ‘False’ CNVs will be assigned 0 point.

##### 3.4 Good-region score

We calculate the accumulative overlap fraction between each CNV and all low-complexity and high repetitive (LCHR) genomic regions, then multiply that value with 100, and use 100 to subtract it, finally we got good-region score.

##### 3.5 Normal score

We calculate the accumulative overlap fraction between each CNV and a list of CNVs with over 1% frequency in normal population, then multiply that value with 100, and use 100 to subtract it, finally we got normal score.

##### 3.6 Pathogenicity score

ClassifyCNV calculates a pathogenicity score for each CNV, depending which the CNV is classified into different pathogenic categories: pathogenic, likely pathogenic, benign, likely benign and uncertain significance. CNVPipe extracts both the pathogenicity score and classification, and integrates them into final output.
